# CD59 Promotes SNARE Complex Assembly via Interaction with the Proline-Rich N-Terminal Domain of VAMP2

**DOI:** 10.64898/2026.08.01.740968

**Authors:** Hui Xiang, Yichen Liu, Jun Feng, Lang Wen

## Abstract

**Objective:** The complement regulatory protein CD59 has been shown to promote SNARE complex assembly, yet its interaction with vesicle-associated membrane protein 2 (VAMP2) remains poorly characterized. This study aims to identify the key domain in VAMP2 that mediates the CD59 interaction and to evaluate whether CD59 point mutations affect SNARE complex assembly.

**Methods:** The interaction between CD59 and VAMP2 was examined by immunofluorescence confocal microscopy and co-immunoprecipitation (co-IP). The effect of CD59 on SNARE complex assembly was assessed by co-expressing CD59 with the three core SNARE proteins (syntaxin-1, SNAP-25, and VAMP2) and detecting complex formation by western blotting. Four CD59 single-point mutants were generated and evaluated in SNARE assembly assays. AlphaFold3 was employed to predict the interaction between CD59 and individual VAMP2 domains (confidence threshold: ipTM + pTM ≥ 0.75). Truncated VAMP2 constructs were further characterized by molecular dynamics simulations and co-IP.

**Results:** (1) CD59 directly bound VAMP2 and promoted SNARE complex assembly without altering individual SNARE protein levels. (2) All four CD59 single-point mutants retained the ability to promote SNARE assembly at a level comparable to wild-type CD59, despite showing differential effects on binding stability in molecular dynamics simulations. (3) The proline-rich (P-rich) N-terminal domain of VAMP2 was identified as the key binding interface; its deletion abolished the CD59 interaction, whereas deletion of the SNARE motif did not.

**Conclusion:** CD59 promotes SNARE complex assembly through interaction with the P-rich N-terminal domain of VAMP2. The examined point mutations do not impair this function, suggesting that these sites may tolerate substitutions or that redundant contact residues maintain the interaction.

## Introduction

Intracellular vesicle trafficking and fusion with target membranes are fundamental to key physiological processes including neurotransmitter release and hormone secretion^[1, 2]^. The critical event in membrane fusion is the recognition and assembly of t-SNAREs (Syntaxin-1A and SNAP25) on the plasma membrane with v-SNAREs (VAMP2) on vesicles through their respective SNARE motifs, forming the SNARE complex whose released free energy drives membrane fusion ^[3, 4]^. Even subtle dysregulation of SNARE complex function can trigger broad pathological consequences^[5]^; for example, amyloid-β oligomers and α-synuclein oligomers disrupt SNARE complex formation by interfering with Syntaxin-1A or VAMP2, respectively^[6]^. Although several regulators such as the Munc18 family have been identified^[7]^, whether additional unknown regulatory elements exist remains an active area of investigation.

The structure of VAMP2, from the N-terminus to the C-terminus, comprises the P-rich NT domain, the SNARE motif, the JMD, and the TMD^[3]^. VAMP2 dysfunction is implicated in various neuropsychiatric disorders^[8, 9]^, and its functional integrity is essential for maintaining normal neurotransmission^[10, 11]^. In the regulation of SNARE complex assembly, Munc18-1, Munc13-1, and Tomosyns constitute a sophisticated regulatory network^[12–14]^. However, it remains unclear whether additional protein factors participate in the regulation of VAMP2 and SNARE complex assembly.

The complement system is a critical component of innate immunity and also serves as a synaptic organizer in the central nervous system^[15, 16]^. Recent studies have revealed that complement components exert “non-canonical” functions, directly modulating glutamatergic neurotransmitter release^[17–20]^. Beyond its classical neuroprotective role in preventing MAC assembly^[21–23]^, the complement regulator CD59 has been shown to bind VAMP2 and promote SNARE complex formation, thereby enhancing synaptic vesicle fusion and neurotransmitter release^[24]^. Similarly, CD59 interacts with VAMP2 and SNAP25 to promote insulin granule exocytosis in pancreatic β-cells^[25]^, and binds syntaxin-3 to trigger Weibel-Palade body exocytosis in endothelial cells^[26]^. However, the specific regions of CD59 and VAMP2 that mediate this interaction, and how these regions affect SNARE complex assembly, remain unknown.

Based on this background, the present study was designed to systematically investigate the structural basis of the CD59-VAMP2 interaction and its impact on SNARE complex assembly. By constructing four single-point mutants of CD59 and four functional domain deletion mutants of VAMP2, combined with AlphaFold3 prediction, molecular dynamics simulations, and co-immunoprecipitation assays, we identified the key domains mediating this interaction. Our findings demonstrate that CD59 promotes SNARE complex assembly by binding to the P-rich NT domain of VAMP2, rather than the SNARE motif. Furthermore, although the four CD59 point mutations differentially affected binding energetics, none of them abolished the assembly-promoting function.

## Materials and methods

### Cell culture and transfection

HEK293T and SH-SY5Y cells were obtained from the laboratory cell bank. Cells were cultured in DMEM high-glucose medium (Meisen Cell) supplemented with 10% fetal bovine serum (Cellbox), 100 U/mL penicillin, and 100 μg/mL streptomycin (Beyotime) at 37℃ in a 5% CO₂ incubator.

One day prior to transfection, cells were seeded into 6-well plates and transfected at 60%–70% confluence with a plasmid-to-reagent ratio of 1:2 (w/v). Cells were harvested 36–48 h post-transfection for subsequent experiments.

### Plasmid construction and site-directed mutagenesis

Human Flag-CD59 (wild-type) and His-VAMP2 (full-length and truncation mutants) plasmids were purchased from Tsingke Biotechnology and Youbao Biotechnology, respectively. Four single-point mutants of CD59 (N18Q, W40E, C64T, N77A) and four domain-deletion mutants of VAMP2 (ΔP-rich NT, ΔSNARE motif, ΔJMD, ΔTMD) were generated using the Fast Mutagenesis System (TransGen Biotech) according to the manufacturer’s instructions. Mutagenic primer sequences are listed in Table S1. All constructs were verified by Sanger sequencing.

### Co-immunoprecipitation and western blotting

Co-immunoprecipitation: HEK293T cells were harvested 48 h post-transfection, washed with ice-cold PBS, and lysed in lysis buffer containing 1% Triton X-100 and PMSF (Solarbio) at 4℃ for 1 h. After centrifugation at 12,000 r/min for 10 min at 4℃, supernatants were collected, and an aliquot was retained as Input. Protein A/G magnetic beads (Tianrenhe) were incubated with 0.5 μg anti-Flag antibody (Proteintech, 66008-4-Ig) and cell lysates overnight at 4℃ with rotation. After magnetic separation and washing, bound proteins were eluted with SDS sample buffer, heated at 95℃ for 10 min, and analyzed by western blotting.

Western blotting: Protein concentrations of cell or tissue lysates were determined by BCA assay (Thermo Fisher Scientific, TL276863). Equal amounts of protein were resolved by SDS-PAGE (12% separating gel) and transferred to PVDF membranes. Membranes were blocked with 5% non-fat milk and incubated with primary antibodies overnight at 4℃, followed by HRP-conjugated secondary antibodies for 1 h at room temperature. Signals were visualized using ECL substrate (Abclonal) on a Tanon-5200 imaging system. Antibodies and their dilutions are listed in Table S2.

### Immunofluorescence

SH-SY5Y cells were seeded onto sterile coverslips in 24-well plates and cultured for 24–48 h to reach 70%–80% confluence. Cells were fixed with 4% paraformaldehyde for 15–20 min at room temperature, permeabilized with 0.3% Triton X-100 for 10–15 min, and blocked with 1% BSA for 1 h. Cells were then incubated with mouse anti-CD59 monoclonal antibody (Proteintech, 68222-1-Ig, 1:400) and rabbit anti-VAMP2 monoclonal antibody (Bioss, bs-1951R, 1:400) overnight at 4℃. After washing, cells were incubated with Alexa Fluor 488-conjugated goat anti-rabbit IgG (1:500) and Alexa Fluor 594-conjugated goat anti-mouse IgG (1:500) for 1 h at room temperature in the dark. Nuclei were stained with DAPI, and coverslips were mounted. Images were acquired using a Leica TCS SP8 confocal microscope.

### AlphaFold3 structure prediction

AlphaFold3 was used to predict the structures of CD59 in complex with full-length VAMP2 and its truncation mutants. Amino acid sequences of target proteins were submitted to the AlphaFold3 server, and the interface predicted Template Modeling score (ipTM) and global predicted Template Modeling score (pTM) were analyzed. The threshold of ipTM + pTM ≥ 0.75, as recommended by Homma et al^[27]^, was used to assess the confidence of protein-protein interaction predictions. Predicted structures were visualized using PyMOL.

### Molecular docking and molecular dynamics simulation

Structure modeling: Homology models of CD59 (wild-type and four mutants) were generated using SWISS-MODEL with PDB ID 2J8B as the template. VAMP2 (full-length and four truncation mutants) were modeled using PDB ID 2KOG as the template. Structures were preprocessed using SPDBV software (energy minimization, disulfide bond formation) and water molecules were removed using PyMOL.

Molecular docking: Rigid-body docking was performed using the HDOCK server, with CD59 as the receptor and VAMP2 as the ligand. Global search docking was conducted, and the conformation with the highest HDOCK score was selected as the initial model for subsequent simulations.

Molecular dynamics simulation: All complex systems were subjected to 100 ns molecular dynamics simulations using the GROMACS package (GROMOS96 54A7 force field). Each system was placed in a periodic water box with counterions added to neutralize the charge. Energy minimization was performed using the steepest descent method, followed by NVT and NPT equilibration (300 K, 1 bar). The simulation step size was 2 fs, and all bonds involving hydrogen atoms were constrained using the LINCS algorithm. Trajectories were saved every 40 ps. Complex stability was evaluated by root-mean-square deviation (RMSD), radius of gyration (Rg), and hydrogen bond number.

### Statistical analysis

Data are expressed as mean ± standard error of the mean (SEM). All experiments were independently repeated at least three times. Statistical analyses were performed using GraphPad Prism 8.0.2. Comparisons between two groups were analyzed using two-tailed unpaired Student’s t-test (with Welch’s correction when variances were unequal). Comparisons among multiple groups were analyzed by one-way ANOVA. Statistical significance was set at p < 0.05.

## Results

### CD59 binds VAMP2 and promotes SNARE complex assembly

Using confocal immunofluorescence microscopy in SH-SY5Y neuroblastoma cells, endogenous CD59 (red) and VAMP2 (green) showed highly overlapping fluorescent signals, indicating that the two proteins are spatially adjacent in cells (Figure 1A). The secondary-antibody control showed no specific staining. To further validate the direct interaction, HEK293T cells were co-transfected with Flag-CD59 and VAMP2, and Co-IP was performed using anti-Flag antibody. Input samples showed successful expression of Flag-CD59 and VAMP2. In the Co-IP fraction, both Flag-CD59 and VAMP2 were detected in the Flag-CD59 transfection group, whereas neither was detected in the Flag-vector control group, confirming that CD59 binds VAMP2 (Figure 1B).

**Figure 1.**
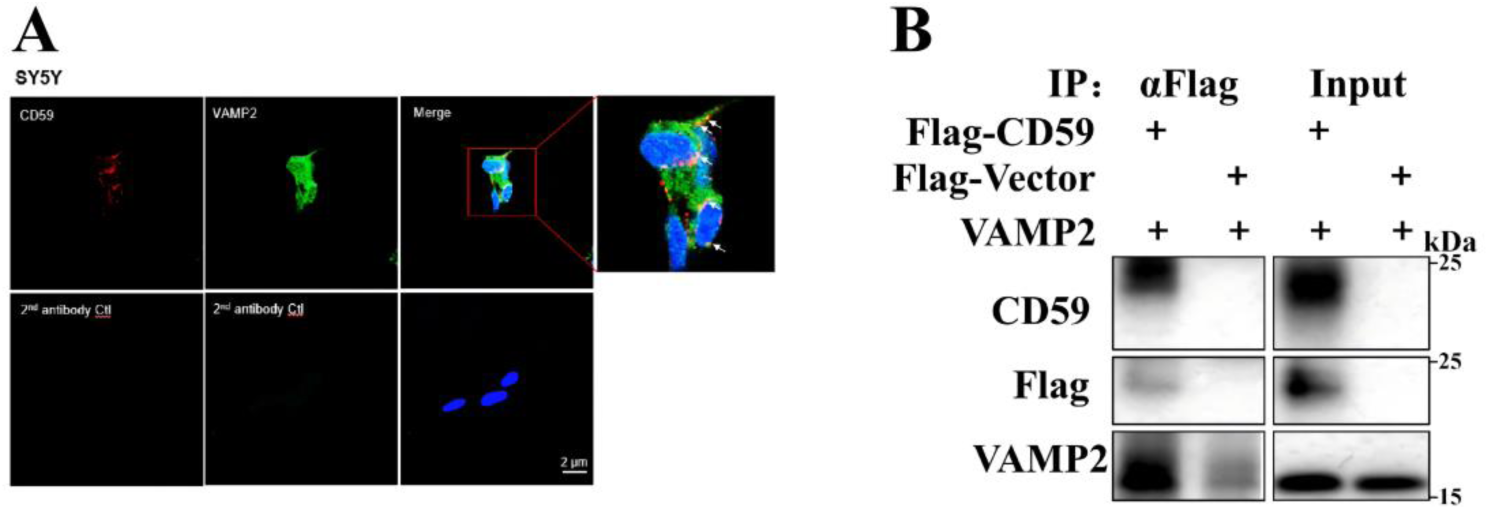
CD59 colocalizes with VAMP2 and interacts with VAMP2 in cells. (A) Confocal immunofluorescence showing endogenous CD59 (red) and VAMP2 (green) in SH-SY5Y cells; nuclei were stained with DAPI (blue). The lower panel shows the secondary-antibody control. Scale bar, 2 μm. (B) Co-IP of Flag-CD59 and VAMP2 in HEK293T cells. Cells were co-transfected with Flag-CD59 or Flag-vector and VAMP2, and anti-Flag immunoprecipitates were analyzed by western blot. Molecular-weight markers (kDa) are indicated on the right.

To investigate the regulatory effect of CD59 on SNARE complex assembly, HEK293T cells were co-transfected with the three core SNARE proteins (VAMP2, Syntaxin-1A, and SNAP25) together with increasing amounts of CD59 plasmid (0, 0.5, or 1 μg). Under non-boiled conditions, the intensity of the high-molecular-weight SNARE complex band increased progressively with CD59 expression (Figure 2A, C). Under boiled conditions, the expression levels of the individual SNARE proteins remained unchanged (Figure 2B, C). These results demonstrate that CD59 promotes SNARE complex assembly without altering the expression levels of individual SNARE proteins.

**Figure 2.**
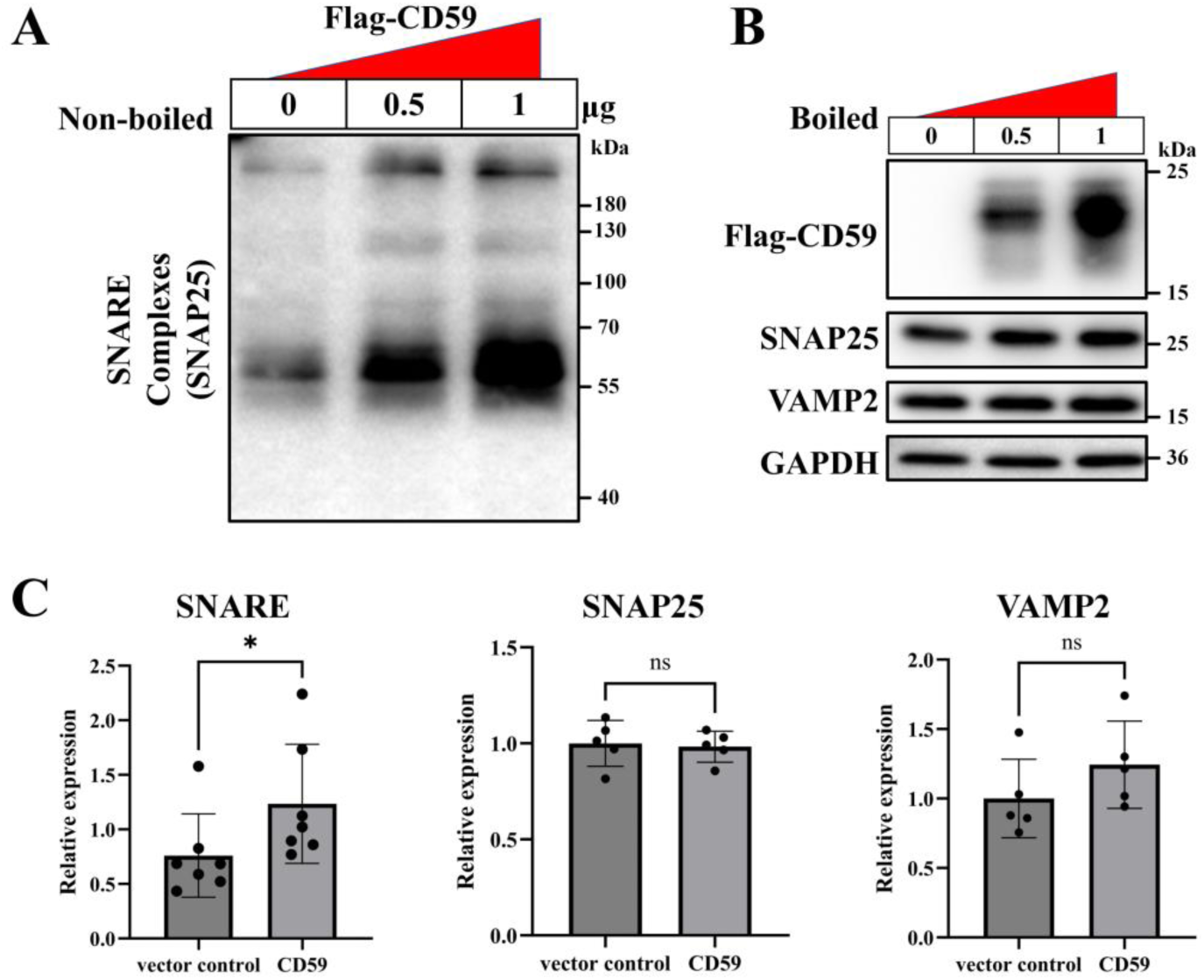
CD59 promotes SNARE complex assembly in a dose-dependent manner without changing SNARE monomer levels. (A) HEK293T cells co-transfected with VAMP2, Syntaxin-1A, and SNAP25 together with increasing amounts of Flag-CD59 (0, 0.5, or 1 μg) were analyzed under non-boiled conditions by western blot using anti-SNAP25 to detect SNARE complex formation. (B) The same cell lysates were boiled and probed for Flag-CD59, SNAP25, VAMP2, and GAPDH. Molecular-weight markers (kDa) are indicated on the right. (C) Quantification of SNARE complex content and monomeric SNAP25 and VAMP2 expression in vector control versus 1 μg CD59 groups. Data are mean ± SEM; ns, not significant; *P < 0.05 versus vector control.

### Construction and validation of CD59 point mutants

Based on the CD59 protein structure (PDB ID 2J8B), human mature CD59 is a flattened ellipsoid of 77 amino acids (Figure 3A). Its secondary structure is dominated by five antiparallel β-strands arranged in a twisted β-sandwich; loops of varying length connect these strands and form functional sites on the molecular surface (Figure 3B). Structural stability depends on five strictly conserved disulfide bonds (Figure 3C), which tightly link the secondary-structure elements and confer a compact, deformation-resistant fold.

**Figure 3.**
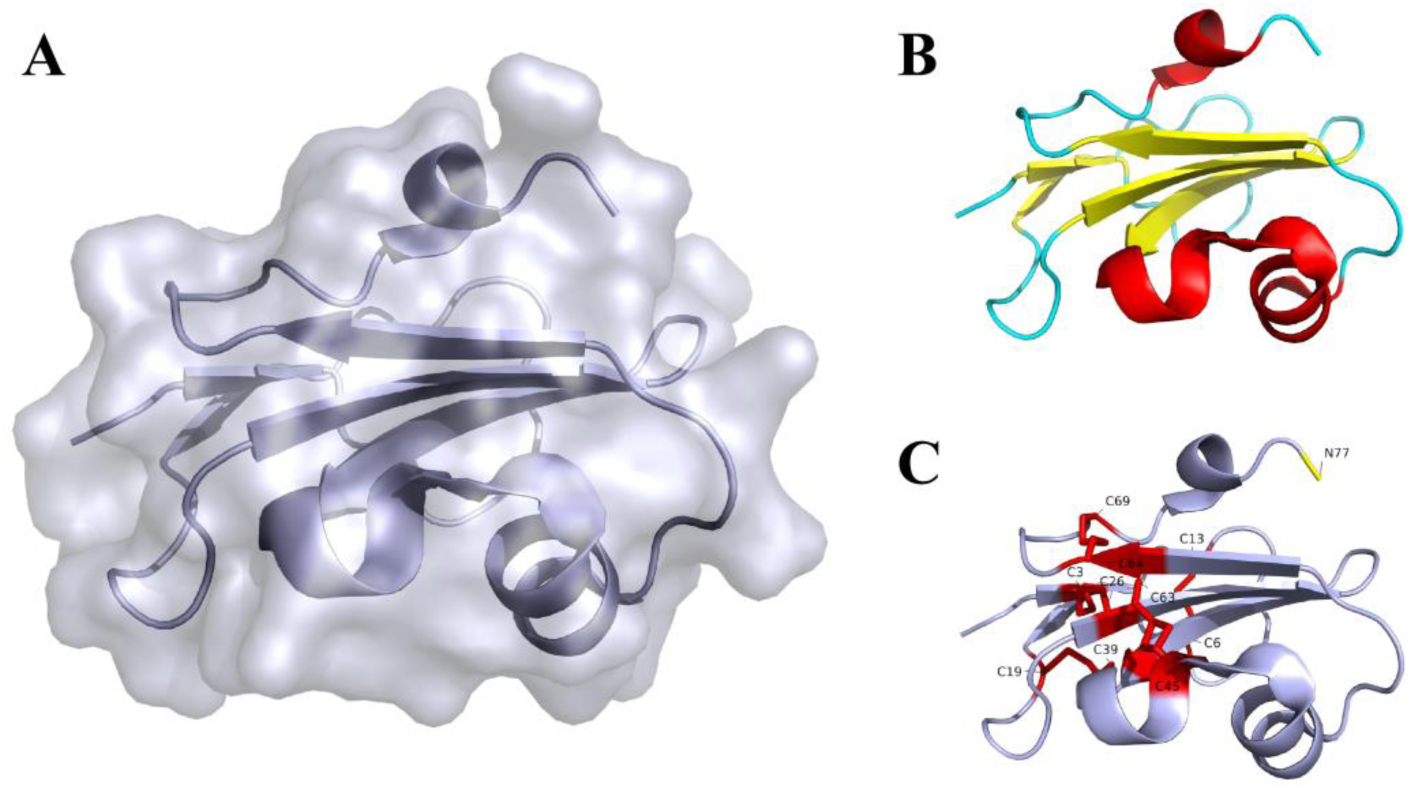
Three-dimensional structure and structural features of CD59. (A) Overlay of the surface representation and polypeptide backbone of CD59. (B) Secondary-structure distribution of CD59; α-helices (red), β-strands (yellow), and random coils (cyan). (C) Distribution of disulfide bonds (red) in CD59.

Four functionally critical residues of human CD59 were selected for single-point mutagenesis (Figure 4A): N18Q (removes N-glycosylation), W40E (abolishes complement inhibitory activity), C64T (a pathogenic mutation associated with CD59 deficiency), and N77A (prevents GPI anchoring). Figure 4B and 4C show the original residue positions and the structures after mutation, respectively; the overall CD59 fold was not markedly altered by any single mutation.

**Figure 4.**
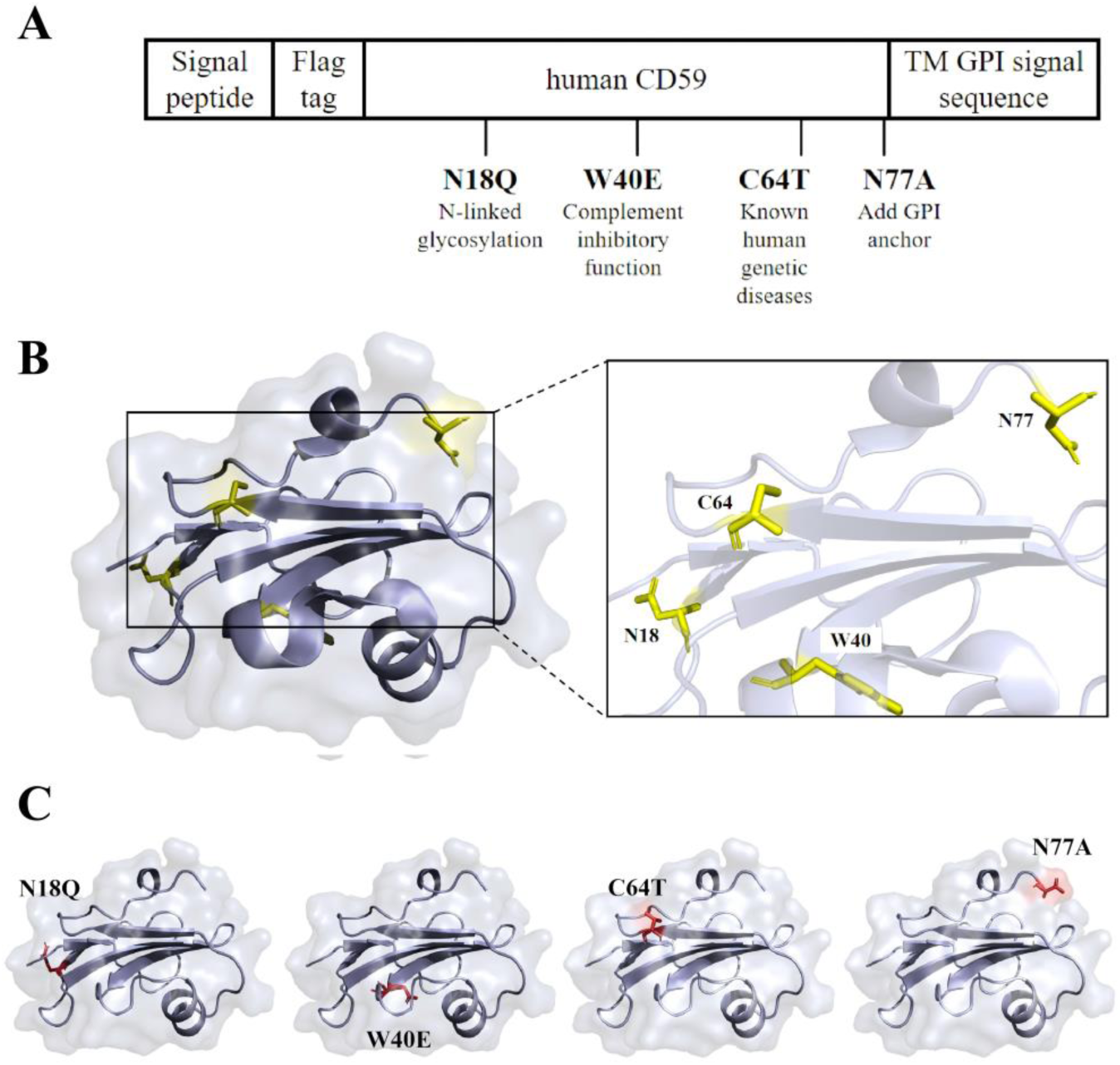
Linear structure and three-dimensional localization of functionally critical CD59 point mutants. (A) Linear schematic of the four CD59 mutants (N18Q, W40E, C64T, and N77A) and their functional annotations. (B) Original amino-acid positions of N18, W40, C64, and N77 in the CD59 structure. (C) Structural models of the N18Q, W40E, C64T, and N77A mutants; mutated residues are highlighted in red.

The Flag-CD59 wild-type and mutant plasmids were verified by Sanger sequencing. Sequence alignment (Figure 5A) and sequencing chromatograms (Figure 5B) showed that the target base substitutions matched the designed mutations without unexpected insertions, deletions, or additional mutations. Western blot analysis showed that, compared with wild-type CD59, the N18Q mutant appeared as a single band with reduced molecular weight, consistent with loss of N-glycosylation (Figure 5C). The W40E, C64T, and N77A mutants showed molecular weights similar to wild-type CD59. For N77A, which blocks GPI-anchor addition, the expressed protein may be present in a soluble secreted form.

**Figure 5.**
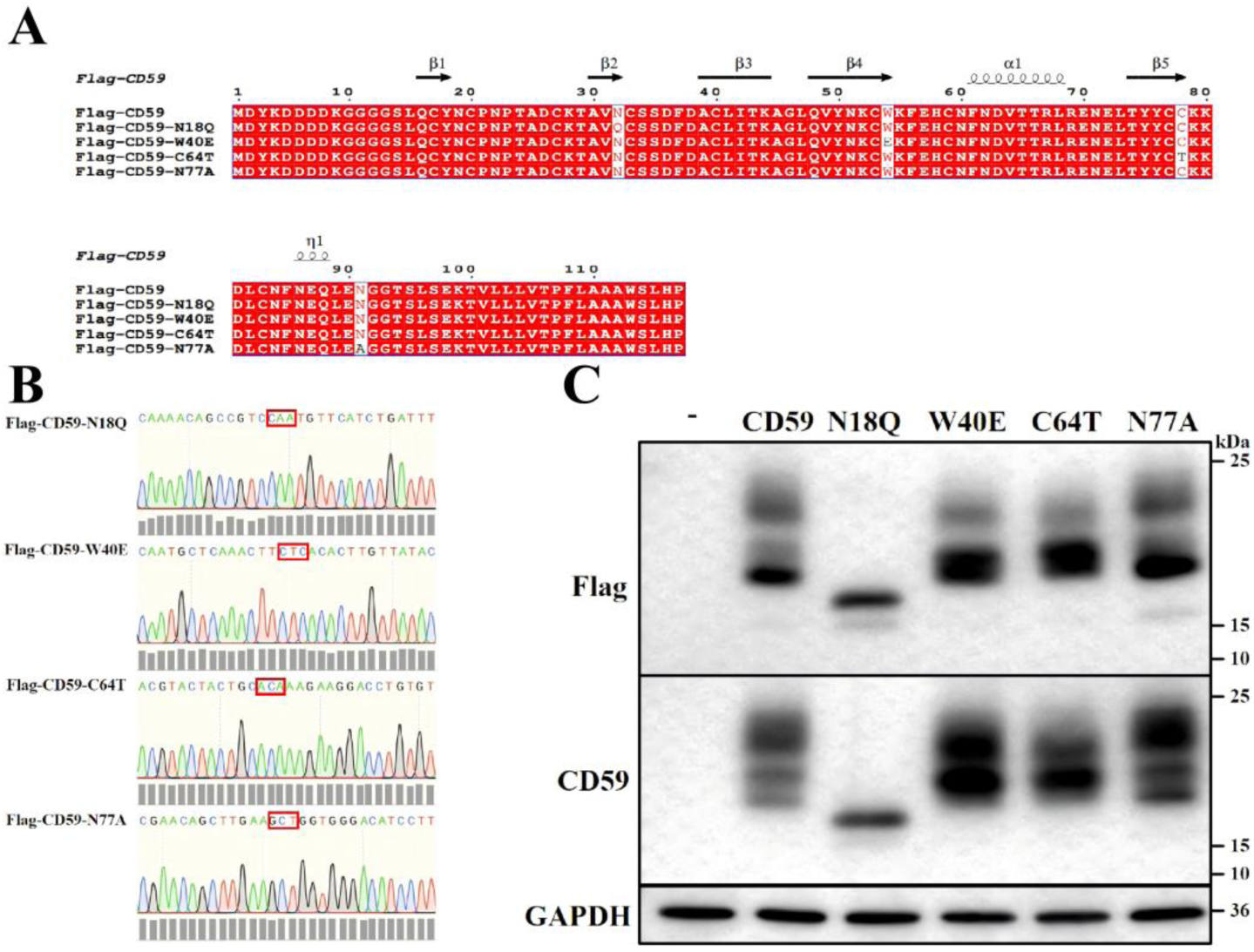
Construction validation and protein expression of CD59 point mutants. (A) Sequence alignment of wild-type (WT) and mutant Flag-CD59 plasmids. (B) Sanger sequencing chromatograms of the target mutation sites (red boxes). (C) Western blot detection of Flag-CD59 WT and mutant proteins using anti-Flag, anti-CD59, and anti-GAPDH antibodies. Molecular-weight markers (kDa) are indicated on the right.

### Molecular dynamics simulations of CD59 and its mutants binding to VAMP2

To interpret the interaction between CD59 and VAMP2, 100 ns molecular dynamics (MD) simulations were performed for CD59 (wild-type and the four mutants) in complex with VAMP2. RMSD, radius of gyration (Rg), and hydrogen-bond number were used to assess complex stability and interaction features. The average RMSD values for CD59, N18Q, W40E, C64T, and N77A bound to VAMP2 were 1.56, 1.58, 1.83, 1.23, and 1.21 nm, respectively (Figure 6A). N18Q and W40E showed increased RMSD relative to wild-type CD59, with W40E showing the largest increase, indicating reduced complex stability. In contrast, C64T and N77A showed decreased RMSD, suggesting a more stable or less fluctuating complex. The Rg values were very similar across all variants (4.29–4.33 nm; Figure 6B), indicating that the mutations had little effect on overall compactness. The average hydrogen-bond numbers were 107, 124, 113, 118, and 122 for CD59, N18Q, W40E, C64T, and N77A, respectively (Figure 6C). All mutants showed slightly more hydrogen bonds than wild-type CD59, with N18Q and N77A showing the largest increases, suggesting that these mutations may strengthen the binding interaction.

**Figure 6.**
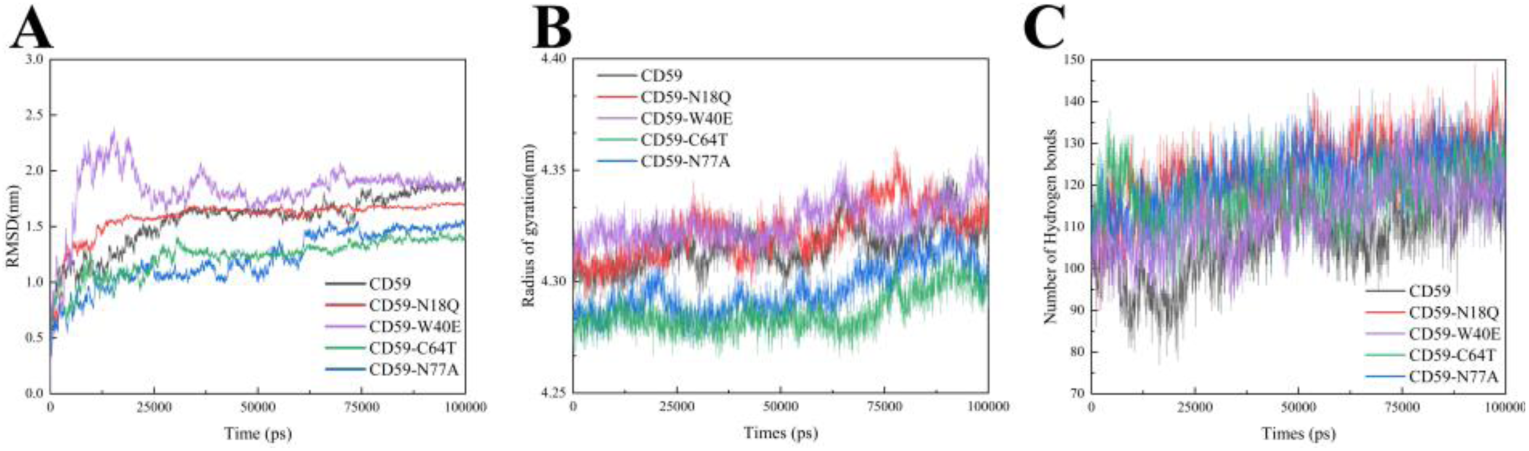
Molecular dynamics simulations of CD59 wild-type and mutants binding to VAMP2. (A) Root-mean-square deviation (RMSD) curves for CD59 WT and mutants (N18Q, W40E, C64T, N77A) in complex with VAMP2. (B) Radius of gyration (Rg) over the 100 ns simulation. (C) Number of hydrogen bonds over the simulation time.

Representative binding models are shown in Figure 7A–E. Compared with wild-type CD59 (Figure 7A), N18Q (Figure 7B) and W40E (Figure 7C) showed no obvious differences in hydrogen-bonding residues, whereas C64T (Figure 7D) and N77A (Figure 7E) formed additional hydrogen-bonding contacts with VAMP2.

**Figure 7.**
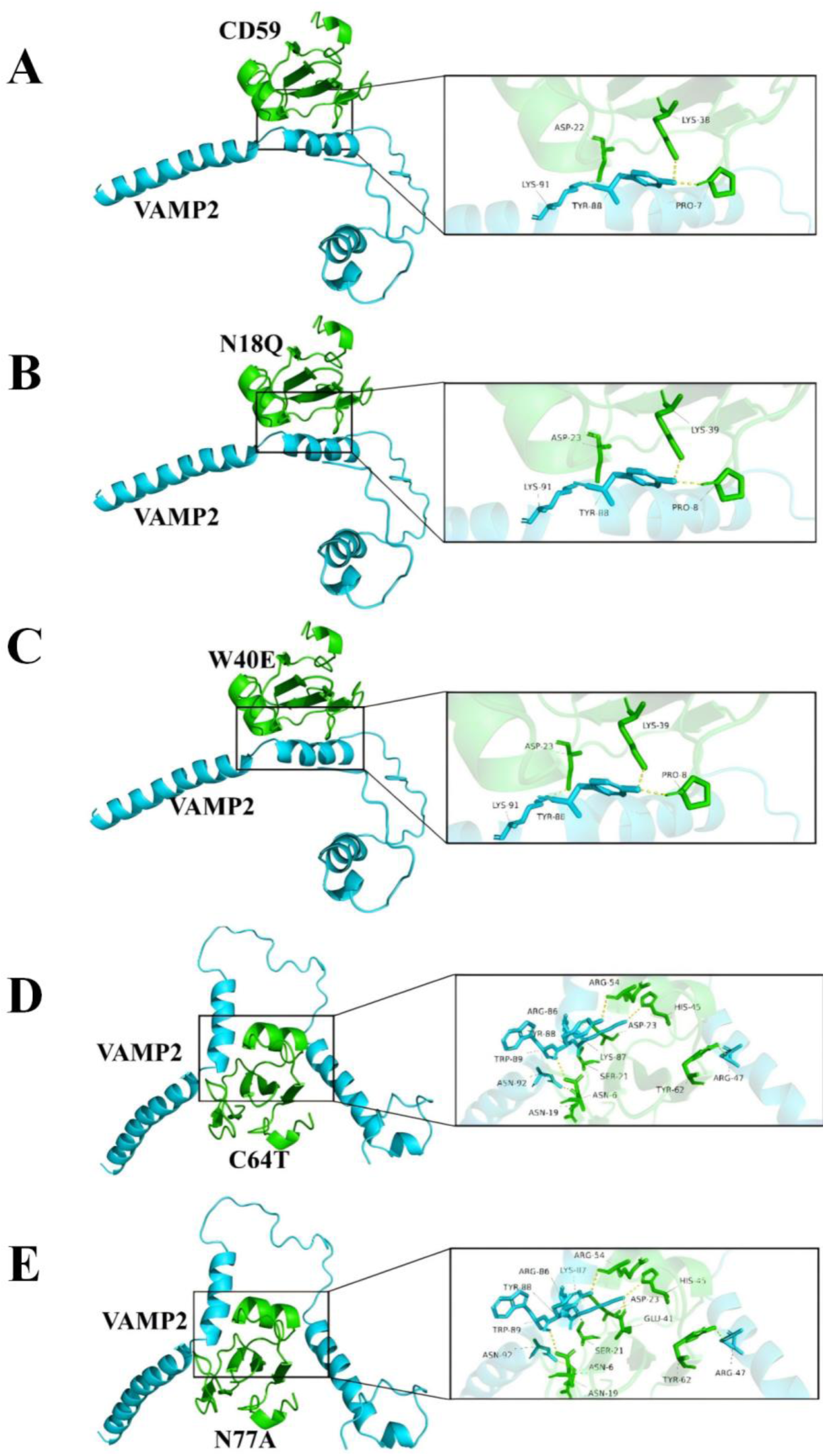
Binding models of CD59 wild-type and mutants with VAMP2. (A–E) Molecular docking structures of wild-type CD59 (A), CD59-N18Q (B), CD59-W40E (C), CD59-C64T (D), and CD59-N77A (E) with VAMP2. VAMP2 is shown in cyan, CD59/mutants in green, and hydrogen bonds as yellow dashed lines.

### Molecular dynamics simulations of CD59 and its mutants binding to the SNARE complex

Similarly, 100 ns MD simulations were performed for CD59 (wild-type and the four mutants) in complex with the pre-assembled SNARE complex. The average RMSD values were 0.38, 0.44, 0.55, 0.45, and 0.59 nm for CD59, N18Q, W40E, C64T, and N77A, respectively (Figure 8A). All mutant complexes showed higher RMSD than the wild-type complex, indicating a general decrease in stability. The Rg values were essentially identical (∼5.01–5.02 nm; Figure 8B), indicating little effect on overall compactness. The average hydrogen-bond numbers were 357, 352, 368, 355, and 359, respectively (Figure 8C); these small differences indicate that the mutations do not substantially alter the binding strength between CD59 and the SNARE complex.

**Figure 8.**
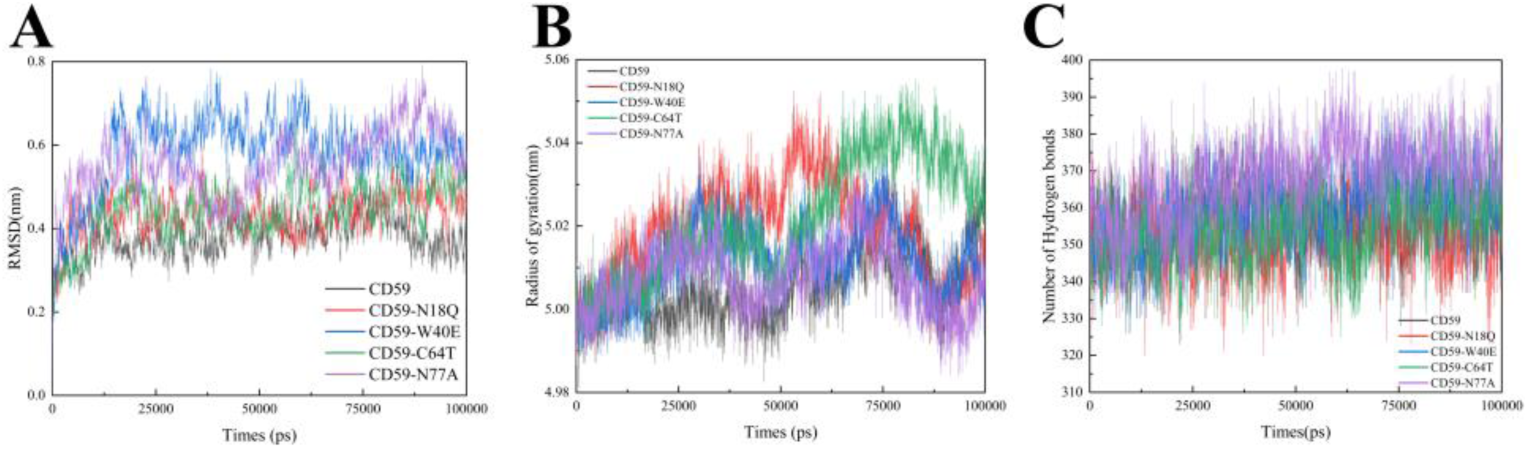
Molecular dynamics simulations of CD59 wild-type and mutants binding to the SNARE complex. (A) RMSD curves for CD59 WT and mutants in complex with the SNARE complex. (B) Radius of gyration over the simulation. (C) Number of hydrogen bonds over the simulation time.

The binding models are shown in Figure 9A–E. Overall, the CD59 mutations did not markedly change the structure of the SNARE complex; hydrogen-bonding sites remained largely stable, with only partial residue changes.

**Figure 9.**
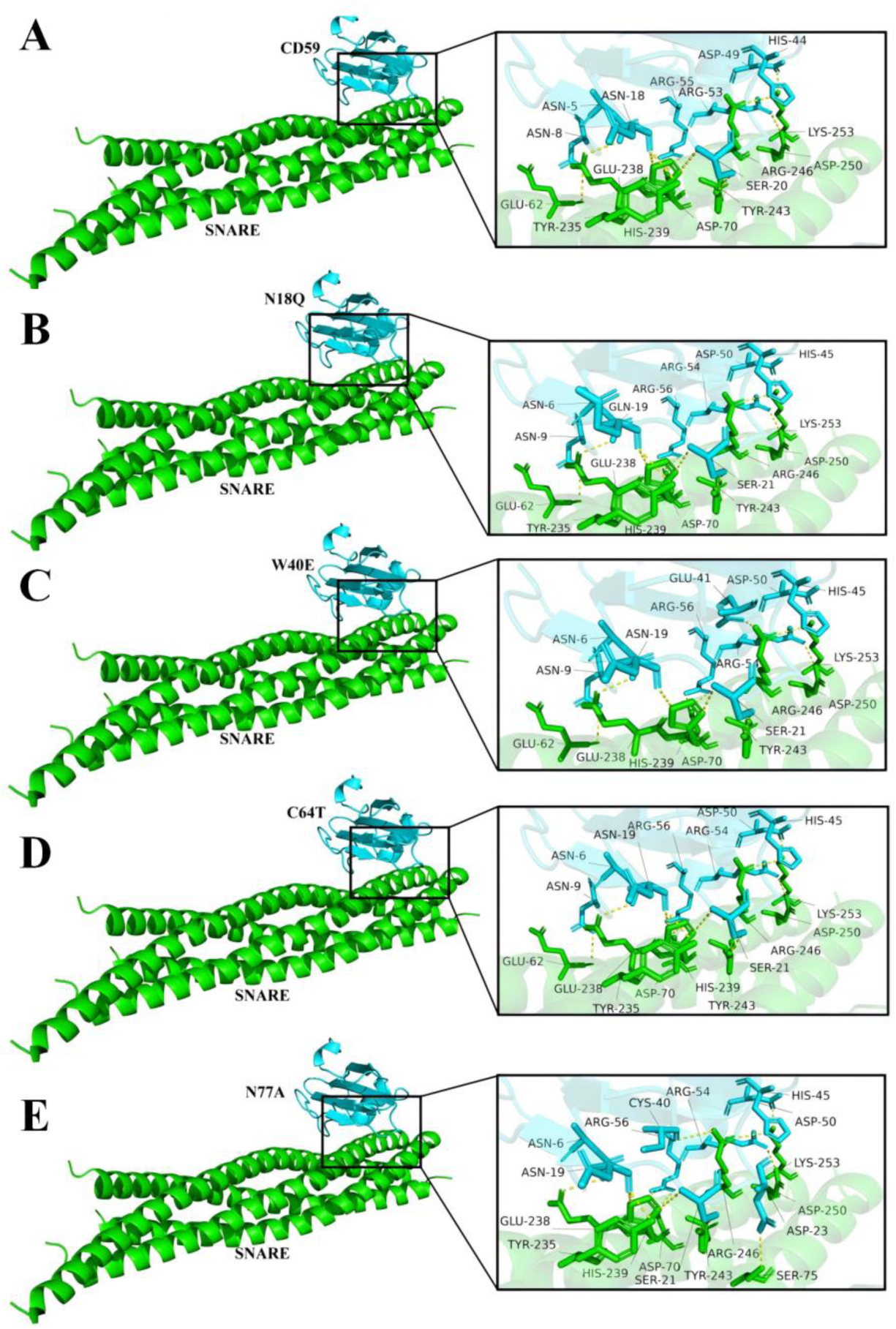
Binding models of CD59 wild-type and mutants with the SNARE complex. (A–E) Molecular docking structures of wild-type CD59 (A), CD59-N18Q (B), CD59-W40E (C), CD59-C64T (D), and CD59-N77A (E) with the SNARE complex. CD59/mutants are shown in cyan, the SNARE complex in green, and hydrogen bonds as yellow dashed lines.

### Effect of CD59 mutants on SNARE complex assembly

We next examined whether the CD59 point mutants retained the ability to promote SNARE complex assembly. Under non-boiled conditions, N18Q, W40E, C64T, and N77A promoted SNARE complex assembly to a similar extent as wild-type CD59, with no significant difference in band intensity (Figure 10A, B). Under boiled conditions, all mutants were expressed normally (Figure 10C), and the expression levels of the core SNARE proteins SNAP25 (Figure 10D) and VAMP2 (Figure 10E) were unchanged compared with the wild-type CD59 group. These results indicate that the four mutations do not impair the ability of CD59 to promote SNARE complex assembly and do not alter SNAP25 or VAMP2 expression levels.

**Figure 10.**
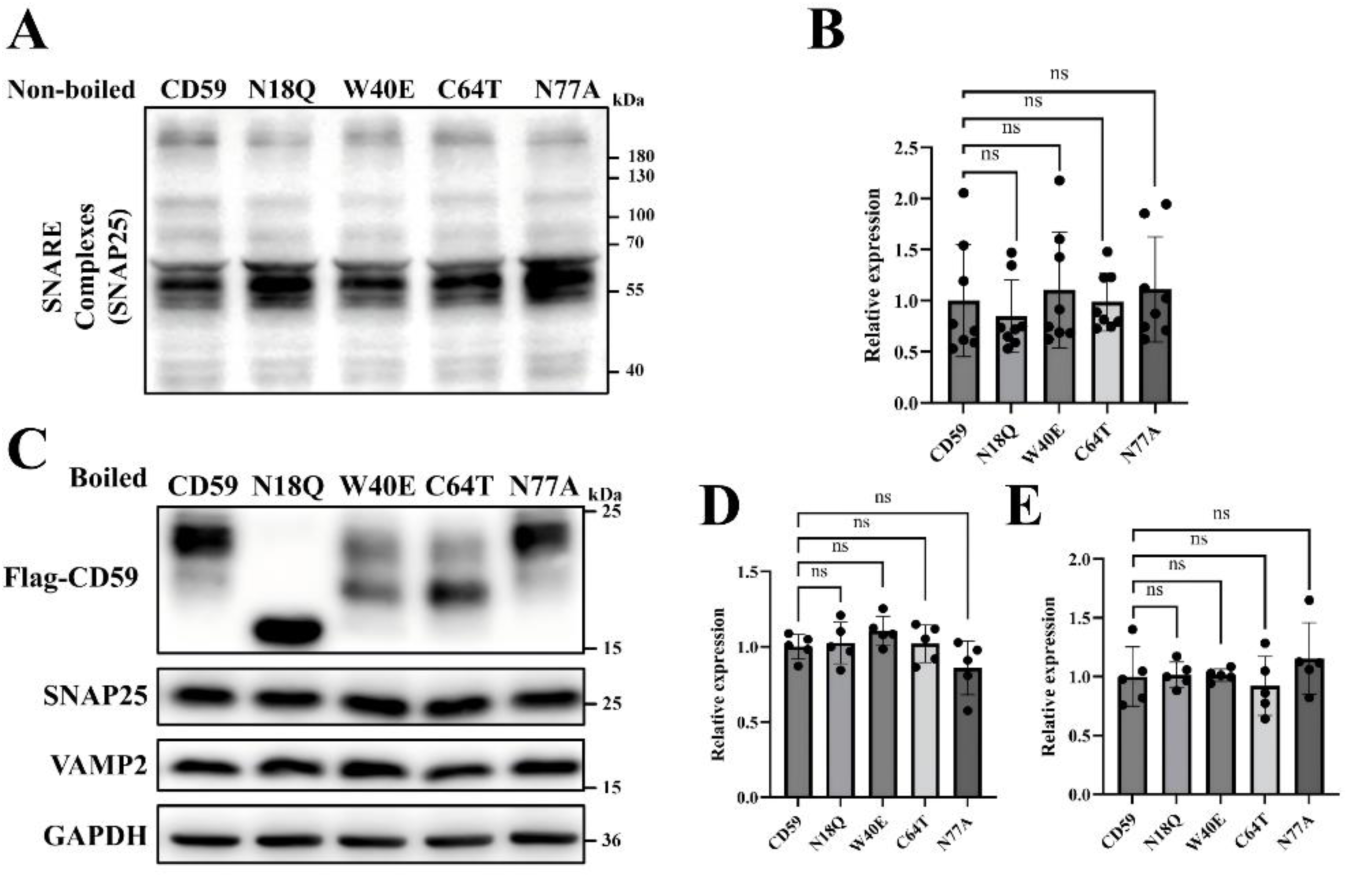
Effect of CD59 mutants on SNARE complex assembly and SNARE protein expression. (A) SNARE complex assembly detected under non-boiled conditions using anti-SNAP25. (B) Quantification of the SNARE complex band in A. (C) Western blot of boiled lysates for Flag-CD59, SNAP25, VAMP2, and GAPDH. (D, E) Quantification of SNAP25 and VAMP2 bands in C. Data are mean ± SEM; ns, not significant.

### AlphaFold3 prediction of VAMP2 domain interactions with CD59

Analysis of the VAMP2 structure (PDB ID 2KOG) showed that VAMP2 is a 116-amino-acid elongated protein composed almost entirely of two long α-helices connected by a flexible loop, with no β-strands (Figure 11A). Based on sequence and function, VAMP2 can be divided into four domains (Figure 11B): the N-terminal proline-rich domain (P-rich NT, residues 1–30), involved in binding interaction factors; the core SNARE motif (residues 31–85), which forms α-helices and assembles into the SNARE complex; the juxtamembrane domain (JMD, residues 86–94), regulating membrane interactions; and the transmembrane domain (TMD, residues 95–116), anchoring VAMP2 to the synaptic vesicle membrane.

**Figure 11.**
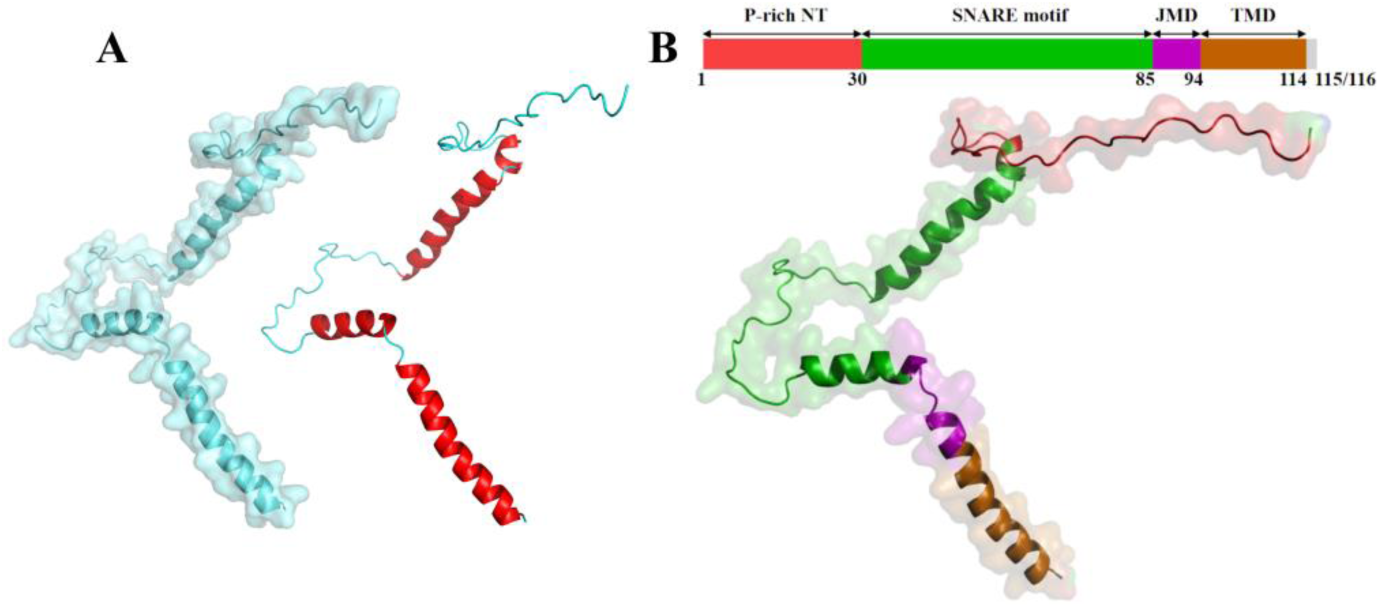
Three-dimensional structure and domain organization of VAMP2. (A) Three-dimensional structural model of VAMP2. (B) Linear domain schematic of VAMP2, showing the P-rich NT, SNARE motif, JMD, and TMD domains.

We submitted the amino-acid sequences of CD59 and VAMP2 to AlphaFold3. The full-length CD59–VAMP2 prediction yielded an ipTM + pTM score of 0.48, below the 0.75 confidence threshold (Figure 12A, B). Visualization showed that AlphaFold3 predicted CD59 interacting mainly with the VAMP2 SNARE motif (Figure 12B), which conflicts with experimental evidence. Previous reports indicate that intrinsically disordered or highly flexible regions in full-length proteins can reduce AlphaFold prediction accuracy[47]; therefore, we used individual VAMP2 domains as inputs instead of the full-length sequence.

**Figure 12.**
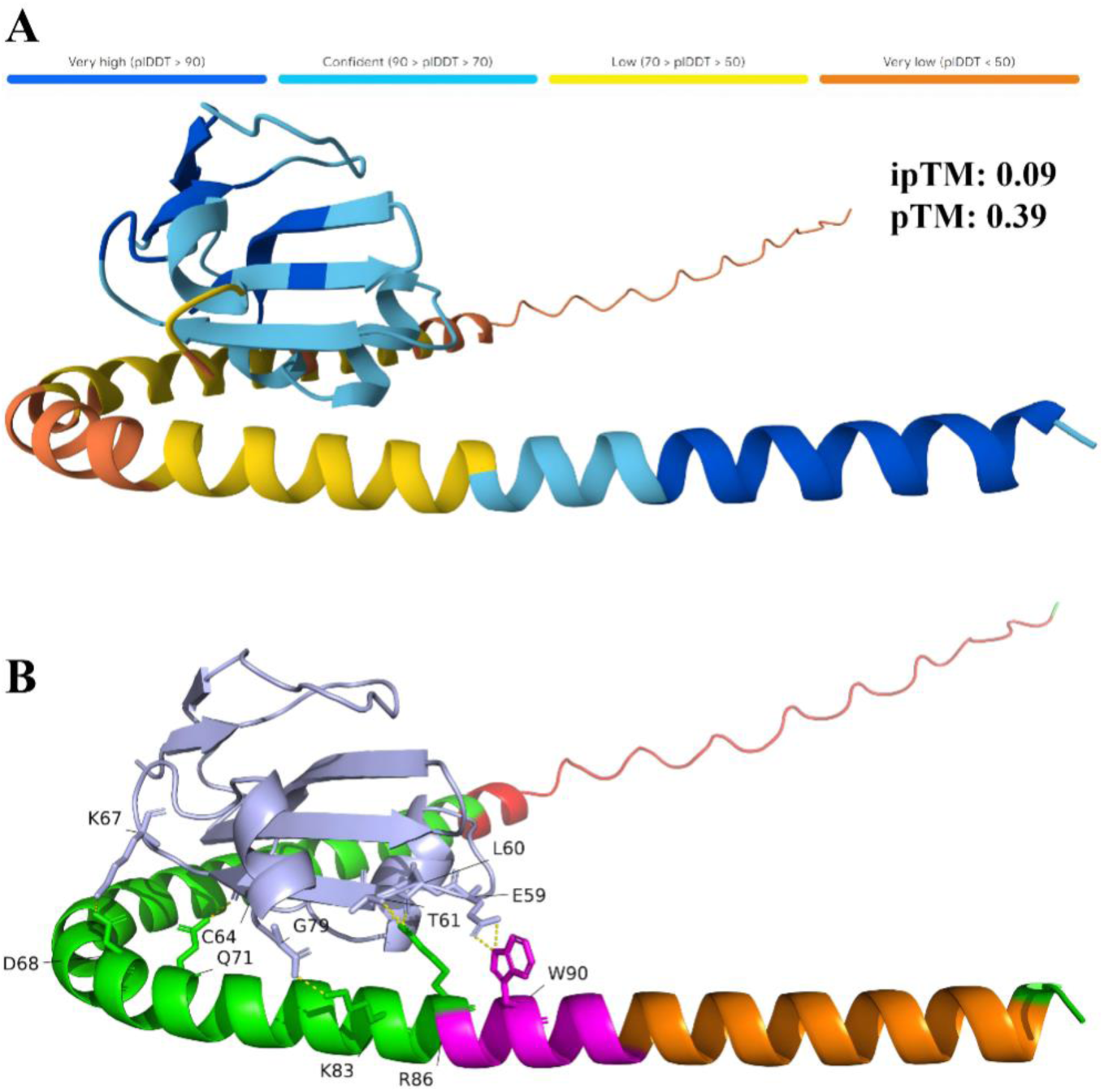
AlphaFold3-predicted interaction model and structural features of full-length VAMP2 with CD59. (A) Overall AlphaFold3 model of the VAMP2–CD59 interaction, colored by pDWT confidence (very high, >90; confident, 70–90; low, 50–70; very low, <50). ipTM = 0.09, pTM = 0.39. (B) Enlarged view of the predicted interaction interface, highlighting potentially interacting residues.

AlphaFold3 was used to predict interactions between full-length CD59 and full-length VAMP2 or each of the four VAMP2 domains (P-rich NT, SNARE motif, JMD, and TMD). The combined ipTM + pTM scores are summarized in Table 1. Full-length VAMP2 gave 0.48, below the confidence threshold, whereas the P-rich NT, JMD, and TMD domains gave scores of 0.85, 1.11, and 1.18, respectively, all above the 0.75 threshold; the SNARE motif gave 0.72, slightly below threshold. High-confidence models for the P-rich NT, JMD, and TMD interactions are shown in Figure 13A, with the interface regions highlighted in Figure 13B. These results suggest that the P-rich NT, JMD, and TMD domains are likely the key regions mediating CD59–VAMP2 interaction.

**Figure 13.**
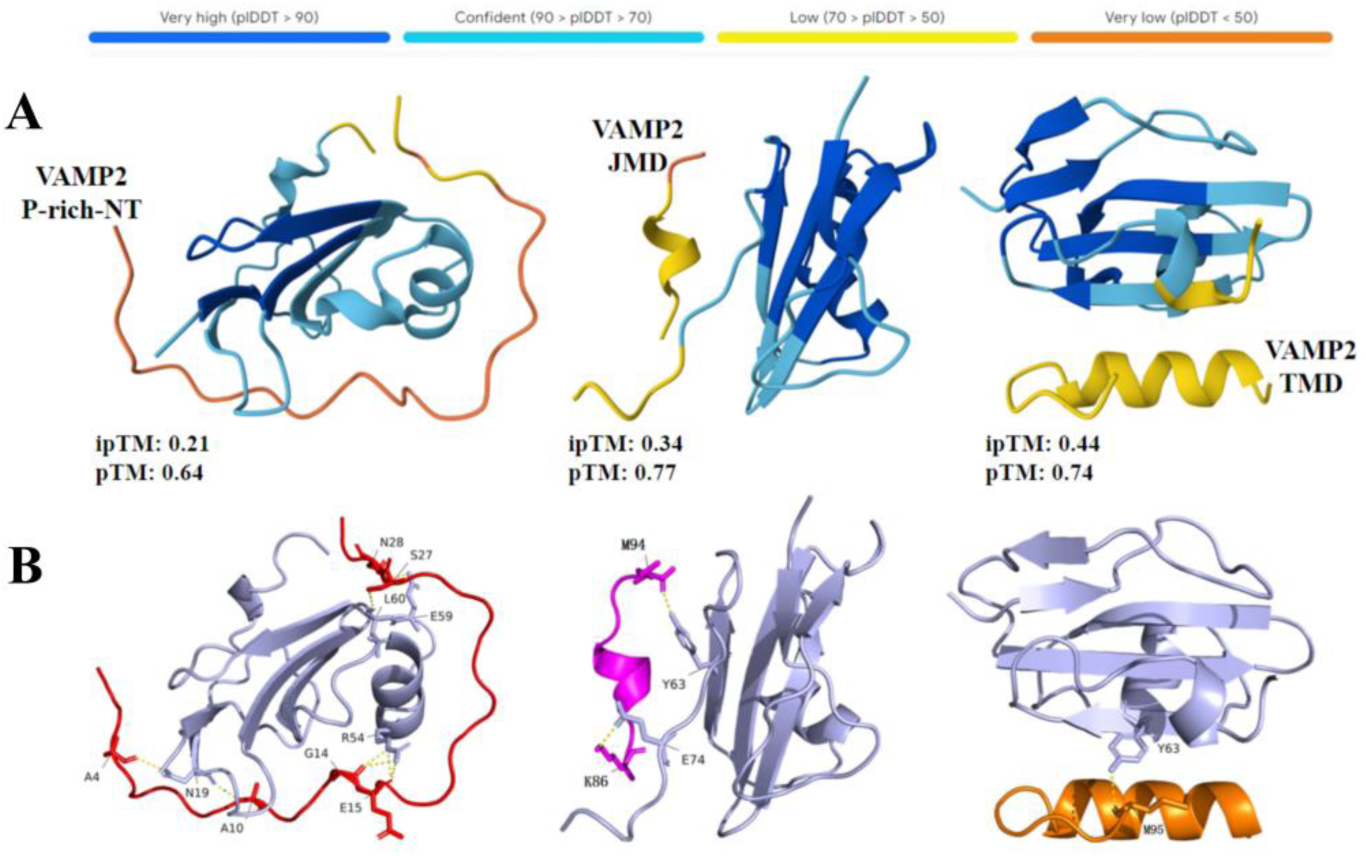
AlphaFold3-predicted interaction models of VAMP2 domains with CD59. (A) Overall models of CD59 with the P-rich NT, JMD, and TMD domains of VAMP2, colored by pDWT confidence; ipTM and pTM scores are indicated. (B) Enlarged interface models showing the spatial contact regions between CD59 and each VAMP2 domain.

**Table 1.** AlphaFold3-predicted interaction scores between CD59 and VAMP2 constructs.

| CD59<br>(PDB ID: 2J8B) | VAMP2<br>(PDB ID: 2KOG) | ipTM | pTM | ipTM+pTM<br>($\geq 0.75$) |
| --- | --- | --- | --- | --- |
| CD59-all | VAMP2-all | 0.09 | 0.39 | 0.48 |
| CD59-all | P-rich-NT | 0.21 | 0.64 | 0.85 |
| CD59-all | SNARE motif | 0.18 | 0.54 | 0.72 |
| CD59-all | JMD | 0.34 | 0.77 | 1.11 |
| CD59-all | TMD | 0.44 | 0.74 | 1.18 |

### Construction of VAMP2 domain-deletion mutants

Based on the P-rich NT, SNARE motif, JMD, and TMD domain architecture of VAMP2, four domain-deletion truncation mutants were designed (Figure 14A). PCR amplification of the truncation constructs produced clear bands of the expected sizes, with no non-specific products (Figure 14B). Sanger sequencing chromatograms were clean and the sequences at the deletion junctions matched the designed truncations (Figure 15A), confirming successful construction of ΔP-rich NT, ΔSNARE motif, ΔJMD, and ΔTMD mutants.

**Figure 14.**
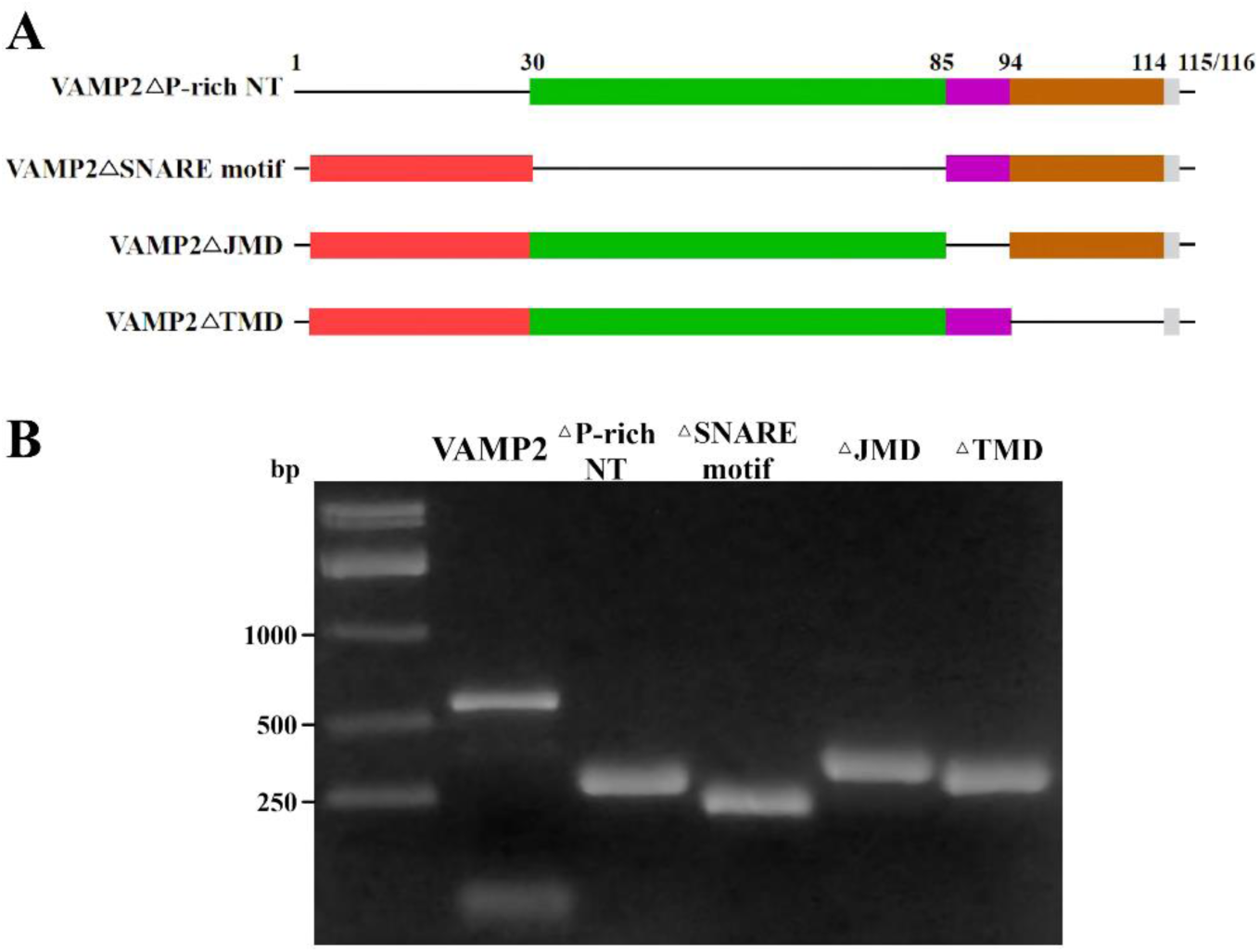
Design and PCR validation of VAMP2 domain-deletion mutants. (A) Linear schematic of the VAMP2 truncation mutants (ΔP-rich NT, ΔSNARE motif, ΔJMD, and ΔTMD), showing the deleted amino-acid ranges. (B) Agarose gel electrophoresis of PCR products for full-length VAMP2 and the truncation mutants. DNA size markers (bp) are shown on the left.

**Figure 15.**
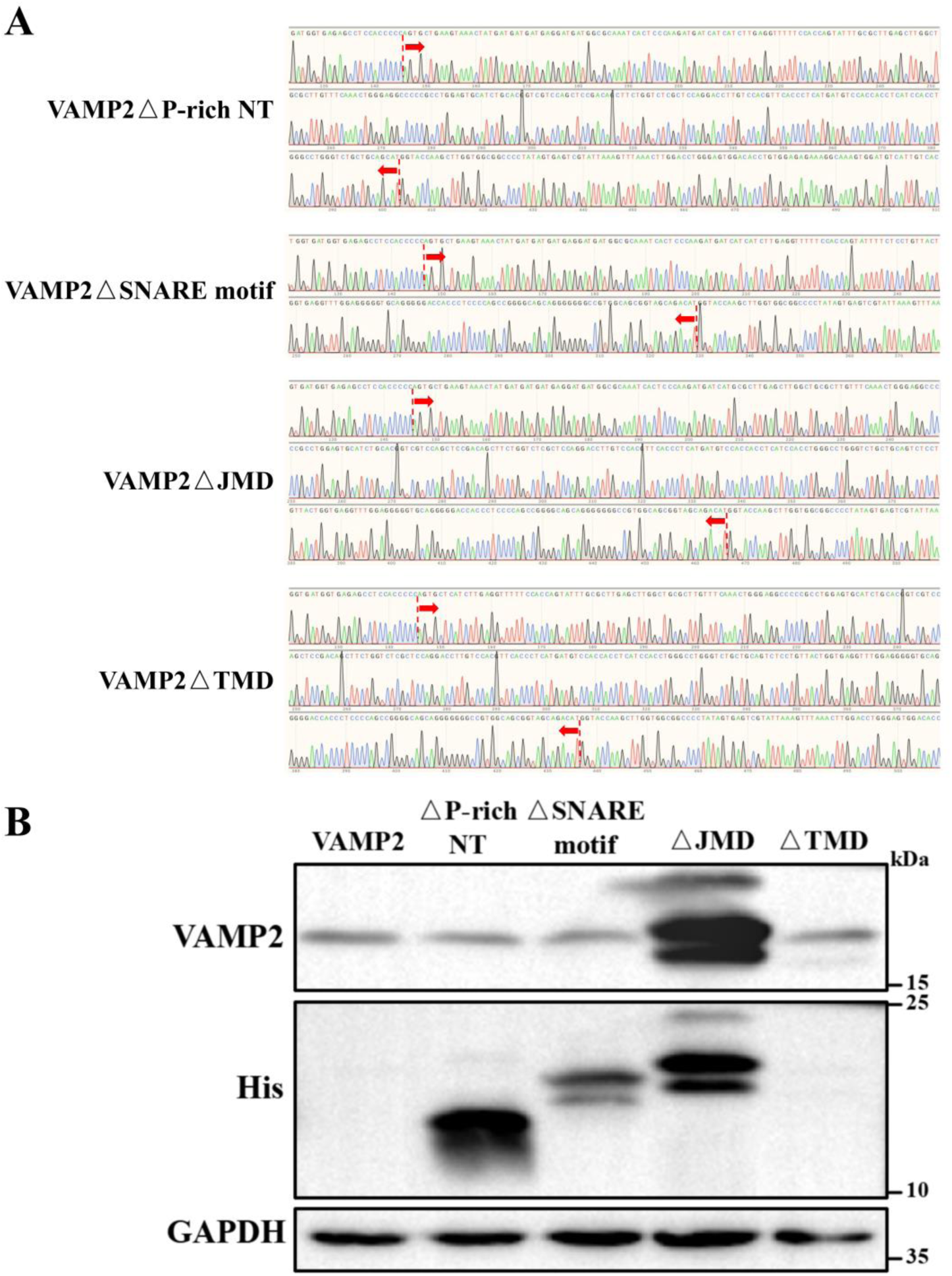
Sequencing validation and protein expression of VAMP2 truncation mutants. (A) Sanger sequencing chromatograms of the truncation junctions (red arrows). (B) Western blot detection of full-length VAMP2 and truncation mutants (ΔP-rich NT, ΔSNARE motif, ΔJMD, and ΔTMD) using anti-VAMP2 and anti-His antibodies.

Western blot analysis showed that VAMP2 bands were detected in all lanes at approximately 15–25 kDa (Figure 15B). His-tagged truncation mutants migrated at different apparent molecular weights depending on the deleted region; the ΔJMD mutant showed the largest molecular weight and highest expression, whereas the ΔP-rich NT mutant showed the smallest molecular weight. GAPDH loading control was consistent across lanes. Together, these results confirm that the VAMP2 truncation mutants were correctly constructed and successfully expressed in cells.

### Molecular dynamics simulations and interaction analysis of VAMP2 truncation mutants with CD59

MD simulations were performed for full-length VAMP2 and the four truncation mutants in complex with CD59. The average RMSD values for VAMP2, ΔP-rich NT, ΔSNARE motif, ΔJMD, and ΔTMD were 1.56, 2.01, 0.78, 1.57, and 1.42 nm, respectively (Figure 16A), indicating that ΔSNARE motif formed the most stable complex and ΔP-rich NT the least stable. The Rg values were 4.32, 4.89, 3.45, 3.86, and 3.79 nm, respectively (Figure 16B), showing that the ΔSNARE motif complex was the most compact and the ΔP-rich NT complex the most extended. Hydrogen-bond analysis showed that ΔSNARE motif formed the fewest hydrogen bonds with CD59, whereas ΔP-rich NT formed the most (Figure 16C).

**Figure 16.**
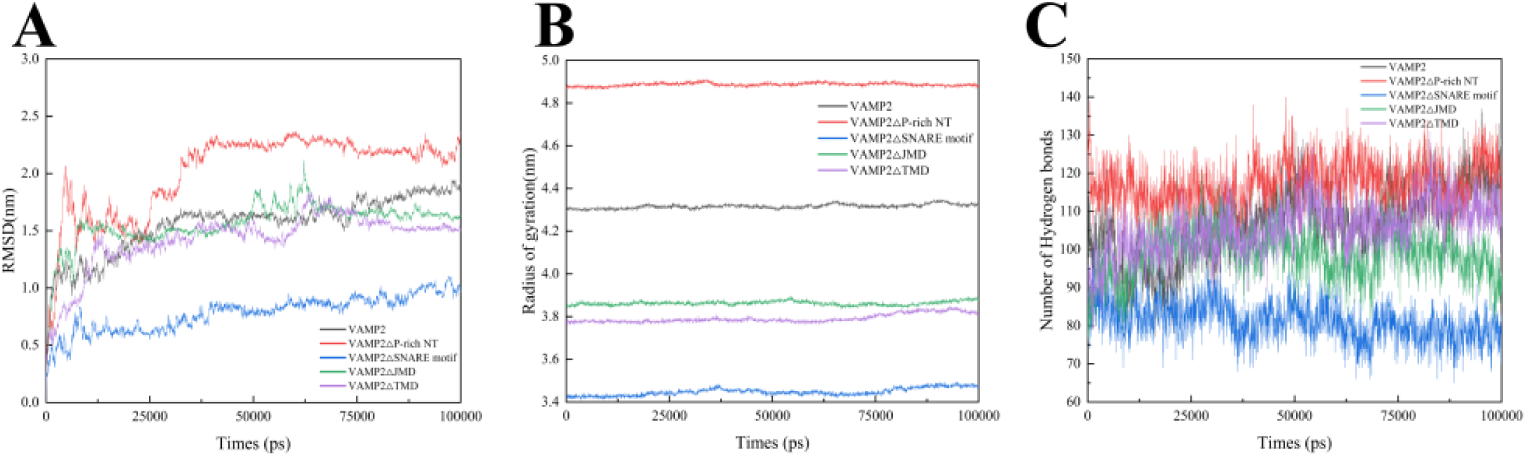
Molecular dynamics simulations of VAMP2 and its truncation mutants binding to CD59. (A) RMSD curves for full-length VAMP2 and truncation mutants (ΔP-rich NT, ΔSNARE motif, ΔJMD, and ΔTMD) in complex with CD59 over 100 ns. (B) Radius of gyration over the simulation. (C) Number of hydrogen bonds over the simulation time.

The binding models are shown in Figure 17A–E. The ΔJMD (Figure 17D) and ΔTMD (Figure 17E) complexes showed stability and hydrogen-bonding patterns comparable to full-length VAMP2 (Figure 17A). The ΔSNARE motif complex (Figure 17C) was the most stable and compact but had the weakest hydrogen-bond interactions. In contrast, the ΔP-rich NT complex (Figure 17B) was the most unstable and extended, with abundant but likely transient hydrogen-bond contacts.

**Figure 17.**
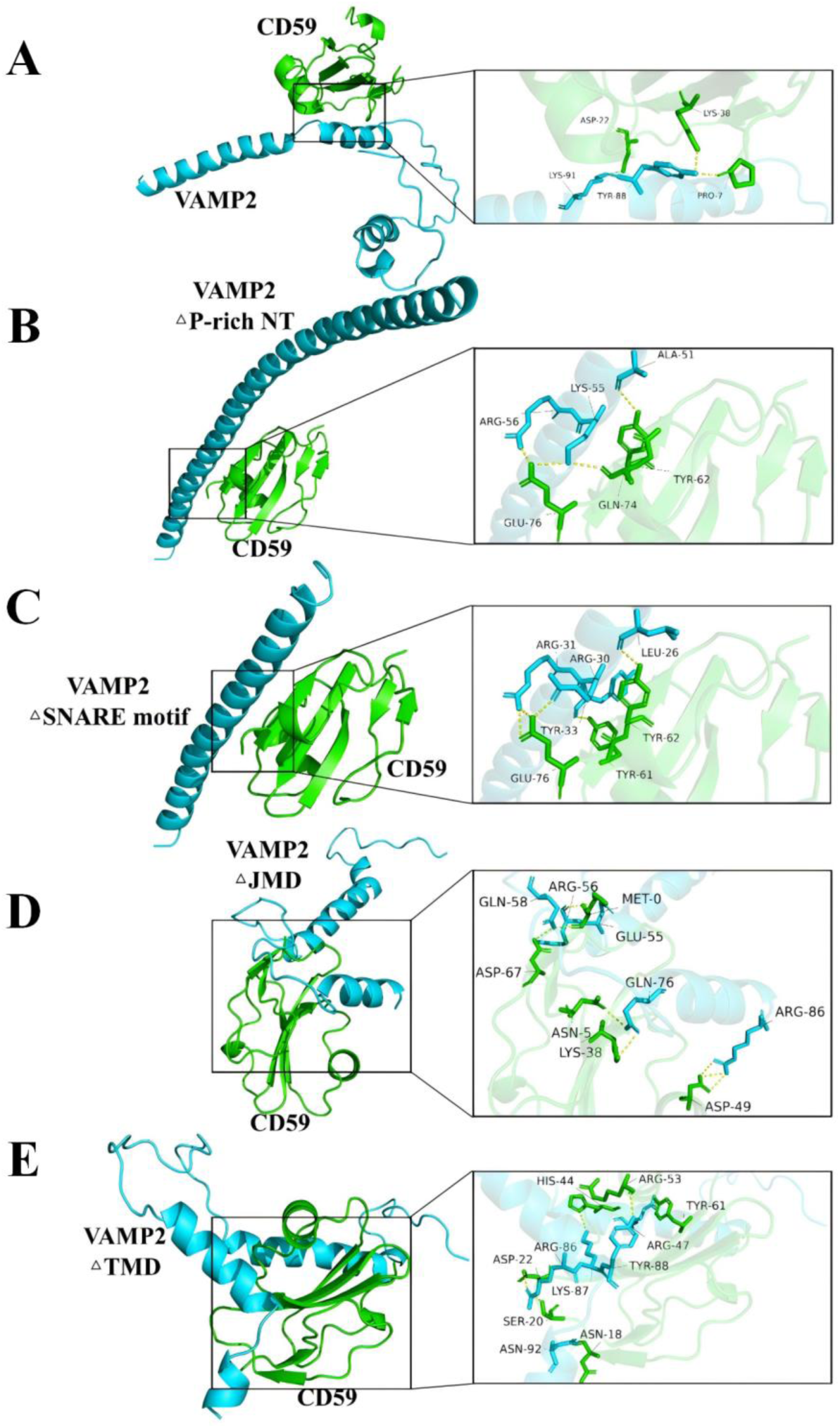
Binding models of VAMP2 and its truncation mutants with CD59. (A–E) Molecular docking structures of full-length VAMP2 (A), VAMP2-ΔP-rich NT (B), VAMP2-ΔSNARE motif (C), VAMP2-ΔJMD (D), and VAMP2-ΔTMD (E) with CD59. CD59 is shown in green, VAMP2 truncations in cyan, and hydrogen bonds as yellow dashed lines.

Co-immunoprecipitation experiments further showed that the ΔSNARE motif truncation still interacted directly with CD59, whereas the ΔP-rich NT truncation did not (Figure 18). These results are consistent with the MD simulations and indicate that the SNARE motif is not the key region for CD59 binding, whereas the P-rich NT domain is essential for the VAMP2–CD59 interaction.

**Figure 18.**
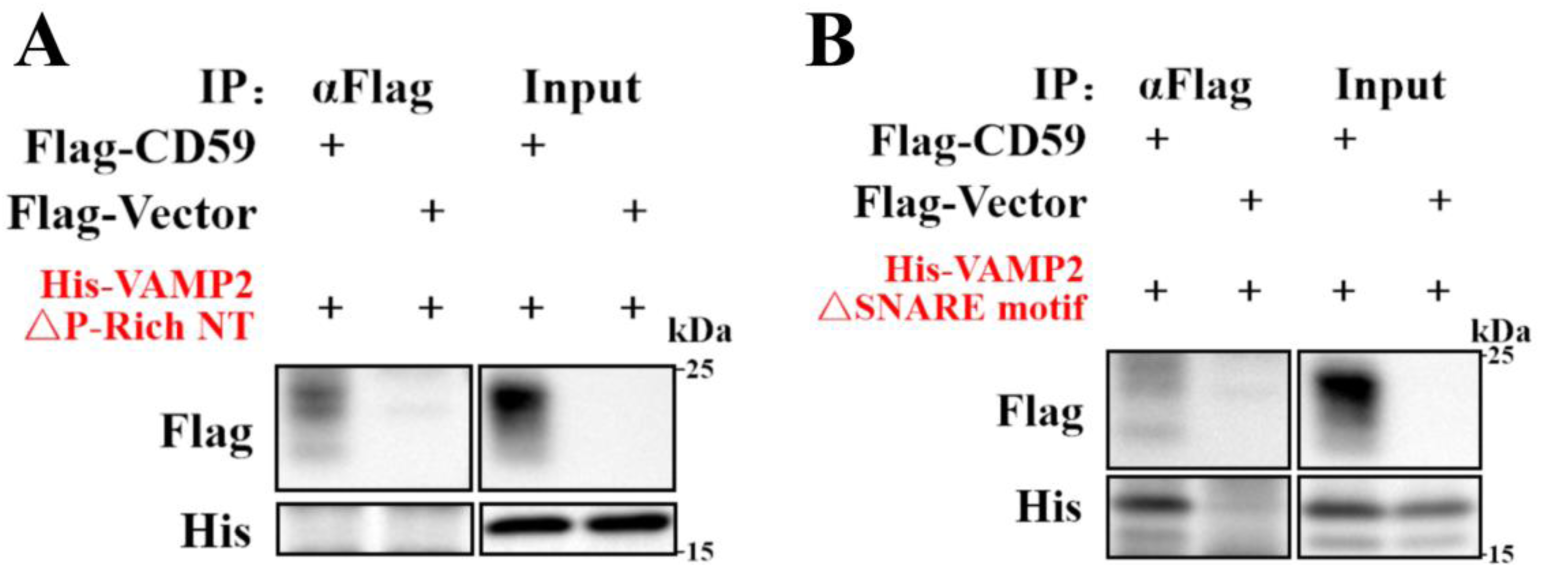
Interaction analysis of VAMP2 ΔP-rich NT and ΔSNARE motif truncation mutants with CD59. (A) HEK293T cells were co-transfected with Flag-CD59 or Flag-vector and His-VAMP2-ΔP-rich NT. (B) HEK293T cells were co-transfected with Flag-CD59 or Flag-vector and His-VAMP2-ΔSNARE motif. Anti-Flag Co-IP was performed and analyzed by western blot. Molecular-weight markers (kDa) are indicated on the right.

## Discussion

In the present study, by constructing four single-point mutants of CD59 and four functional domain deletion mutants of VAMP2, we systematically investigated the mechanism by which CD59 regulates SNARE complex assembly. Our results demonstrate that CD59 promotes SNARE complex assembly, and this effect does not depend on alterations in the expression levels of individual SNARE proteins such as SNAP25 and VAMP2. Through co-immunoprecipitation assays, we confirmed a direct interaction between CD59 and VAMP2. Although molecular dynamics simulations suggested that this interaction is differentially affected by residues N18, W40, C64, and N77, cell-based overexpression experiments showed that none of these mutations significantly impaired the ability of CD59 to promote SNARE complex assembly. Combined with AlphaFold3 structure prediction and molecular dynamics simulations, we found that the P-rich NT domain of VAMP2 is likely the key region mediating its interaction with CD59, whereas the SNARE motif is not involved in this interaction.

The precise assembly of the SNARE complex is a core event in intracellular vesicle fusion, and this process is controlled by multiple regulatory factors^[12, 28–30]^. CD59 has traditionally been regarded as a GPI-anchored complement regulatory protein that protects host cells by inhibiting membrane attack complex formation^[22, 23]^., research on whether CD59 regulates SNARE complex assembly has been primarily advanced by the research group led by Ewelina Golec. They demonstrated that the neuron-specific CD59 isoforms IRIS-1/2 directly bind to the vesicular SNARE protein VAMP2, promote its interaction with target membrane SNAREs, and stabilize SNARE complex assembly^[31]^. Similarly, our study also shows that CD59 promotes SNARE complex formation. More importantly, control experiments confirmed that CD59 overexpression did not alter the total expression levels of individual SNARE proteins such as SNAP25 and VAMP2, ruling out the possibility of indirect promotion of assembly through upregulation of substrate protein expression.

Dissecting the key functional sites of CD59 can reveal the structural basis of the CD59-VAMP2 interaction. Previous studies have shown that the N18Q mutation removes N-glycosylation of CD59, but the mutant protein can still be correctly folded and expressed on the cell surface, indicating that N-glycosylation is not involved in the complement inhibitory activity of CD59^[32, 33]^; the W40E mutation completely abolishes the complement inhibitory activity of CD59^[32]^; the C64T mutation disrupts a disulfide bond in CD59^[34, 35]^; and the N77A mutation eliminates the GPI anchor of CD59^[36]^. In the present study, we found that wild-type CD59 appeared as a diffuse band, indicating varying degrees of glycosylation modification under normal conditions, whereas the N18Q mutation removed N-glycosylation and the corresponding band appeared as a relatively homogeneous single band. Further investigation revealed of the CD59-VAMP2 interaction, we found that the N18Q, W40E, C64T, and N77A mutations all affected the binding between CD59 and VAMP2 to varying degrees. Unexpectedly, however, these mutants still exhibited similar promoting effects on SNARE complex assembly compared with wild-type CD59. This result differs somewhat from the findings of Golec et al^[37]^—although our N18Q mutant showed impaired binding, it still promoted SNARE complex assembly. This discrepancy may arise from significant sequence differences between human and rat CD59 near the GPI anchor region^[37]^. Alternatively, we speculate that, as a GPI-anchored protein, CD59 is enriched in lipid rafts^[38]^. Even if certain mutations weaken the direct binding between CD59 and VAMP2, CD59 may still indirectly promote SNARE complex formation by maintaining lipid raft structural integrity or recruiting other fusion-related factors.

Our results demonstrate that CD59 binds to the proline-rich N-terminus, the juxtamembrane domain, and the transmembrane domain of VAMP2, but not to the SNARE motif that forms the core helical bundle. This binding mode ensures that CD59 does not occupy the core interface required for SNARE zippering, spatially allowing it to accompany the assembly process as a chaperone-like protein. Existing evidence has confirmed that CD59 further participates in the subsequent regulation of membrane fusion: Krus et al. observed that CD59 silencing reduced the proportion of events with fusion pore flickering and accelerated vesicle emptying^[39]^; Gao et al. found that intermittent hypoxia promoted the binding of internalized CD59 to syntaxin-3, increasing calcium influx and promoting Weibel-Palade body exocytosis^[26]^. Combined with our findings, it can be speculated that CD59, anchored near the membrane, resides at the fusion interface and may either reduce the energy barrier of the fusion pore by modulating membrane curvature or undergo allosteric regulation with other SNARE components, collectively influencing fusion pore behavior. Thus, CD59 likely plays multiple roles: acting as a dynamic chaperone to promote SNARE assembly in the early stage, and remaining engaged in the initiation and expansion of the fusion pore after assembly is complete.

Of note, although AlphaFold has demonstrated outstanding performance in structure prediction, it may still yield low-confidence scores for some protein-protein interactions that have been experimentally validated^[40–42]^. In this study, co-immunoprecipitation confirmed a specific interaction between CD59 and VAMP2, but when we submitted full-length VAMP2 to AlphaFold3 for prediction, the resulting ipTM + pTM score was only 0.48, far below the generally accepted high-confidence threshold. The discrepancy between prediction and experimental results is consistent with a functional interaction. Yin et al. found that AlphaFold predictions for antibody-antigen complexes almost entirely failed^[43]^; Pereira et al. also pointed out that the model has a systematic bias toward small-interface complexes^[44]^. The conformational flexibility of disordered regions in full-length proteins reduces AlphaFold prediction accuracy^[45]^. Therefore, in this study, we used the four individual domains of VAMP2 as inputs for prediction and applied the ipTM + pTM ≥ 0.75 threshold recommended by Homma et al^[27]^ to assess confidence. Our results indicate that AlphaFold3 predicts that the interaction between CD59 and VAMP2 likely occurs through the P-rich NT, JMD, and TMD domains.

VAMP2, as a key v-SNARE protein, performs its functions through different structural domains [4]. To investigate how different domains of VAMP2 affect its interaction with CD59, we performed 100 ns molecular dynamics simulations on the full-length protein and four truncation mutants. We found that the binding modes of the SNARE motif and the P-rich NT are completely different. The ΔSNARE motif truncation mutant exhibited the smallest conformational fluctuation (RMSD = 0.78 nm) and the most compact structure (Rg = 3.45 nm), but formed the fewest hydrogen bonds with CD59. This “rigid structure with sparse hydrogen bonds” feature is consistent with previous reports on the conformational plasticity of the SNARE motif^[14, 46]^. In contrast, the ΔP-rich NT truncation mutant showed the largest conformational fluctuation (RMSD = 2.01 nm) and the least compact structure (Rg = 4.89 nm), but formed the densest hydrogen bond network with CD59. This “dynamic but non-specific” binding mode closely resembles the fuzzy complex stage of intrinsically disordered proteins during their initial binding with chaperones^[47]^.

This study preliminarily reveals the roles of key functional sites of CD59 and domains of VAMP2 in their interaction, but several important questions remain to be addressed. For example, the dynamic interaction mechanism between CD59 and VAMP2, and the influence of CD59 localization in lipid raft microenvironments, require further confirmation through techniques such as live-cell imaging and single-molecule approaches. In addition, AlphaFold3 predictions suggest that CD59 may interact with the P-rich NT, JMD, and TMD domains of VAMP2; future studies could employ structural biology methods such as NMR or cryo-electron microscopy to further resolve the complex structure.

In summary, our study demonstrates that CD59 promotes SNARE complex assembly by targeting the P-rich NT domain of VAMP2, rather than the SNARE motif. Although the four point mutations of CD59 differentially affected its binding to VAMP2, none of them abolished the assembly-promoting function, suggesting that this regulatory mechanism possesses functional robustness. This study provides a structural basis for understanding the non-canonical synaptic functions of CD59 and offers a potential theoretical foundation for future intervention strategies targeting SNARE assembly regulation.

## Supplementary Information

**Table S1.**
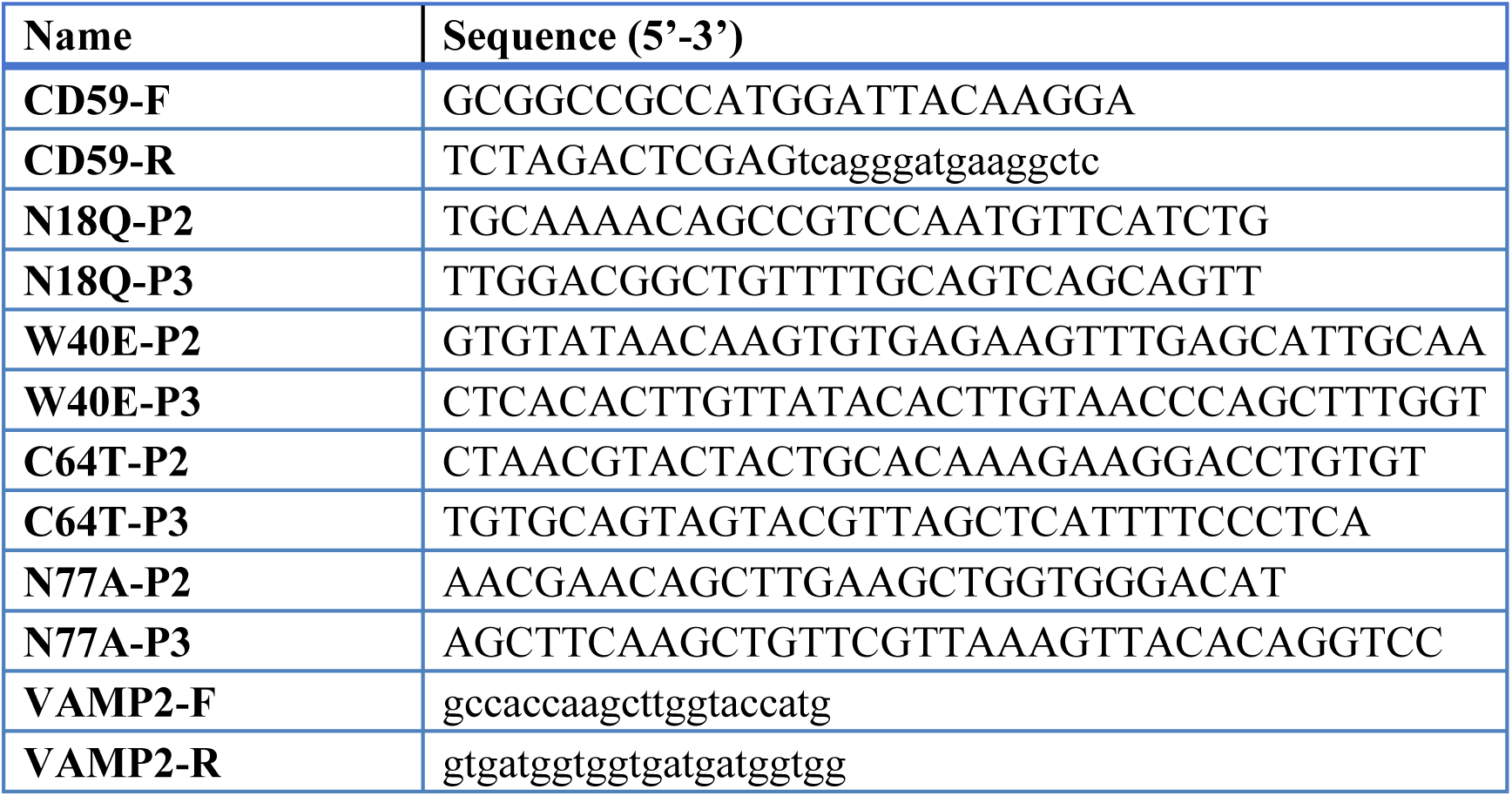
Primer sequences used in this study.

**Table S2.** Antibodies used for western blotting and their dilution ratios.

| Antibody | Dilution | Source | Catalog No. |
| --- | --- | --- | --- |
| Rabbit anti-GAPDH | 1:10000 | Servicebio | GB15004 |
| Rabbit anti-VAMP2 | 1:2500 | MCE | HY-P81673 |
| Mouse anti-Flag tag | 1:10000 | Proteintech | 66008-4-Ig |
| Mouse anti-CD59 | 1:10000 | Proteintech | 68222-1-Ig |
| Mouse anti-SNAP25 | 1:1000 | Biolegend | 836304 |
| Rabbit anti-His tag | 1:5000 | Bioss | 33004R |
| Rabbit anti-VAMP2 (IF) | 1:400 | Bioss | bs-1951R |

## Notes

### Competing Interest Statement

The authors have declared no competing interest.

### Funding

- Hunan Provincial Natural Science Foundation of China: 2025JJ60167
- Hunan college students' Innovation Training Program: 2025437

## References

[1] Cui L, Li H, Xi Y, et al. Vesicle trafficking and vesicle fusion: mechanisms, biological functions, and their implications for potential disease therapy [J]. Mol Biomed, 2022, 3(1): 29.

[2] Stanton A E, Hughson F M. The machinery of vesicle fusion [J]. Curr Opin Cell Biol, 2023, 83: 102191.

[3] Yan C, Jiang J, Yang Y, et al. The function of VAMP2 in mediating membrane fusion: An overview [J]. Front Mol Neurosci, 2022, 15: 948160.

[4] Chen F, Chen H, Chen Y, et al. Dysfunction of the SNARE complex in neurological and psychiatric disorders [J]. Pharmacol Res, 2021, 165: 105469.

[5] Vadisiute A, Meijer E, Szabó F, et al. The role of snare proteins in cortical development [J]. Dev Neurobiol, 2022, 82(6): 457–75.

[6] Margiotta A. Role of SNAREs in Neurodegenerative Diseases [J]. Cells, 2021, 10(5).

[7] André T, Van Berkel A A, Singh G, et al. Reduced Protein Stability of 11 Pathogenic Missense STXBP1/MUNC18-1 Variants and Improved Disease Prediction [J]. Biol Psychiatry, 2024, 96(2): 125–36.

[8] Yoo G, Yeou S, Son J B, et al. Cooperative inhibition of SNARE-mediated vesicle fusion by α-synuclein monomers and oligomers [J]. Sci Rep, 2021, 11(1): 10955.

[9] Ponterio G, Faustini G, El Atiallah I, et al. Alpha-Synuclein is Involved in DYT1 Dystonia Striatal Synaptic Dysfunction [J]. Mov Disord, 2022, 37(5): 949–61.

[10] Rothman J E, Grushin K, Bera M, et al. Turbocharging synaptic transmission [J]. FEBS Lett, 2023, 597(18): 2233–49.

[11] Hoogstraaten R I, Van Keimpema L, Toonen R F, et al. Tetanus insensitive VAMP2 differentially restores synaptic and dense core vesicle fusion in tetanus neurotoxin treated neurons [J]. Sci Rep, 2020, 10(1): 10913.

[12] Hao X, Zhu B, Yang P, et al. SNAP25 mutation disrupts metabolic homeostasis, steroid hormone production and central neurobehavior [J]. Biochim Biophys Acta Mol Basis Dis, 2021, 1868(2): 166304.

[13] Kalyana Sundaram R V, Jin H, Li F, et al. Munc13 binds and recruits SNAP25 to chaperone SNARE complex assembly [J]. FEBS Lett, 2020, 595(3): 297–309.

[14] Meijer M, Öttl M, Yang J, et al. Tomosyns attenuate SNARE assembly and synaptic depression by binding to VAMP2-containing template complexes [J]. Nat Commun, 2024, 15(1): 2652.

[15] Carpanini S M, Torvell M, Bevan R J, et al. Terminal complement pathway activation drives synaptic loss in Alzheimer’s disease models [J]. Acta Neuropathol Commun, 2022, 10(1): 99.

[16] Schartz N D, Tenner A J. The good, the bad, and the opportunities of the complement system in neurodegenerative disease [J]. J Neuroinflammation, 2020, 17(1): 354.

[17] Pittaluga A, Torre V, Olivero G, et al. Non-canonical Roles of Complement in the CNS: From Synaptic Organizer to Presynaptic Modulator of Glutamate Transmission [J]. Curr Neuropharmacol, 2025, 23(7): 820–34.

[18] Benoit M E, Tenner A J. Complement protein C1q-mediated neuroprotection is correlated with regulation of neuronal gene and microRNA expression [J]. J Neurosci, 2011, 31(9): 3459–69.

[19] Perez-Alcazar M, Daborg J, Stokowska A, et al. Altered cognitive performance and synaptic function in the hippocampus of mice lacking C3 [J]. Exp Neurol, 2013, 253: 154–64.

[20] Merega E, Di Prisco S, Lanfranco M, et al. Complement selectively elicits glutamate release from nerve endings in different regions of mammal central nervous system [J]. J Neurochem, 2014, 129(3): 473–83.

[21] Ziabska K, Ziemka-Nalecz M, Pawelec P, et al. Aberrant Complement System Activation in Neurological Disorders [J]. Int J Mol Sci, 2021, 22(9).

[22] Couves E C, Gardner S, Voisin T B, et al. Structural basis for membrane attack complex inhibition by CD59 [J]. Nat Commun, 2023, 14(1): 890.

[23] Patel B, Silwal A, Eltokhy M A, et al. Deciphering CD59: Unveiling Its Role in Immune Microenvironment and Prognostic Significance [J]. Cancers (Basel), 2024, 16(21).

[24] Wen L, Yang X, Wu Z, et al. The complement inhibitor CD59 is required for GABAergic synaptic transmission in the dentate gyrus [J]. Cell Rep, 2023, 42(4): 112349.

[25] Golec E, Ekström A, Noga M, et al. Alternative splicing encodes functional intracellular CD59 isoforms that mediate insulin secretion and are down-regulated in diabetic islets [J]. Proc Natl Acad Sci U S A, 2022, 119(24): e2120083119.

[26] Gao S, Emin M, Thoma T, et al. Complement promotes endothelial von Willebrand factor and angiopoietin-2 release in obstructive sleep apnea [J]. Sleep, 2021, 44(4).

[27] Homma F, Huang J, Van Der Hoorn R A L. AlphaFold-Multimer predicts cross-kingdom interactions at the plant-pathogen interface [J]. Nat Commun, 2023, 14(1): 6040.

[28] Abramian A, Hoogstraaten R I, Murphy F H, et al. Rabphilin-3a negatively regulates neuropeptide release, through its SNAP25 interaction [J]. Elife, 2024, 13.

[29] Bera M, Ramakrishnan S, Coleman J, et al. Molecular determinants of complexin clamping and activation function [J]. Elife, 2022, 11.

[30] Bose D, Bera M, Norman C A, et al. Minimal presynaptic protein machinery governing diverse kinetics of calcium-evoked neurotransmitter release [J]. Nat Commun, 2024, 15(1): 10741.

[31] Golec E, Olsson R, Tuysuz E C, et al. Neuronal CD59 isoforms IRIS-1 and IRIS-2 as regulators of neurotransmitter release with implications for Alzheimer’s disease [J]. Alzheimers Res Ther, 2025, 17(1): 11.

[32] Bodian D L, Davis S J, Morgan B P, et al. Mutational analysis of the active site and antibody epitopes of the complement-inhibitory glycoprotein, CD59 [J]. J Exp Med, 1997, 185(3): 507–16.

[33] Rudd P M, Morgan B P, Wormald M R, et al. The glycosylation of the complement regulatory protein, human erythrocyte CD59 [J]. J Biol Chem, 1997, 272(11): 7229–44.

[34] Nevo Y, Ben-Zeev B, Tabib A, et al. CD59 deficiency is associated with chronic hemolysis and childhood relapsing immune-mediated polyneuropathy [J]. Blood, 2012, 121(1): 129–35.

[35] Karbian N, Eshed-Eisenbach Y, Tabib A, et al. Molecular pathogenesis of human CD59 deficiency [J]. Neurol Genet, 2018, 4(6): e280.

[36] Rushmere N K, Harrison R A, Van Den Berg C W, et al. Molecular cloning of the rat analogue of human CD59: structural comparison with human CD59 and identification of a putative active site [J]. Biochem J, 1994, 304 (Pt 2): 595–601.

[37] Golec E, Rosberg R, Zhang E, et al. A cryptic non-GPI-anchored cytosolic isoform of CD59 controls insulin exocytosis in pancreatic β-cells by interaction with SNARE proteins [J]. FASEB J, 2019, 33(11): 12425–34.

[38] Voisin T B, Couves E C, Tate E W, et al. Dynamics and Molecular Interactions of GPI-Anchored CD59 [J]. Toxins (Basel), 2023, 15(7).

[39] Krus U, King B C, Nagaraj V, et al. The complement inhibitor CD59 regulates insulin secretion by modulating exocytotic events [J]. Cell Metab, 2014, 19(5): 883–90.

[40] Schmid E W, Walter J C. Predictomes, a classifier-curated database of AlphaFold-modeled protein-protein interactions [J]. Mol Cell, 2025, 85(6).

[41] Manyilov V D, Ilyinsky N S, Nesterov S V, et al. Chaotic aging: intrinsically disordered proteins in aging-related processes [J]. Cell Mol Life Sci, 2023, 80(9): 269.

[42] Chojnowski G. gapTrick-structural characterization of protein-protein interactions using AlphaFold [J]. Bioinformatics, 2025, 41(9).

[43] Yin R, Feng B Y, Varshney A, et al. Benchmarking AlphaFold for protein complex modeling reveals accuracy determinants [J]. Protein Sci, 2022, 31(8): e4379.

[44] Pereira G P, Gouzien C, Souza P C T, et al. Challenges in predicting Protac-mediated protein-protein interfaces with AlphaFold reveal a general limitation on small interfaces [J]. Bioinform Adv, 2025, 5(1): vbaf056.

[45] Lin P-Y, Huang S-C, Chen K-L, et al. Analysing protein complexes in plant science: insights and limitation with AlphaFold 3 [J]. Bot Stud, 2025, 66(1): 14.

[46] Salpietro V, Malintan N T, Llano-Rivas I, et al. Mutations in the Neuronal Vesicular SNARE VAMP2 Affect Synaptic Membrane Fusion and Impair Human Neurodevelopment [J]. Am J Hum Genet, 2019, 104(4): 721–30.

[47] Morris O M, Torpey J H, Isaacson R L. Intrinsically disordered proteins: modes of binding with emphasis on disordered domains [J]. Open Biol, 2021, 11(10): 210222.

